# Idioblasts accumulating anticancer alkaloids in *Catharanthus roseus* leaves are a unique cell type

**DOI:** 10.1101/2023.02.24.529939

**Authors:** Joana G. Guedes, Rogério Ribeiro, Inês Carqueijeiro, Ana Luísa Guimarães, Cláudia Bispo, John Archer, Herlander Azevedo, Nuno A. Fonseca, Mariana Sottomayor

## Abstract

*Catharanthus roseus* leaves produce a range of monoterpenoid indole alkaloids (MIAs) that include low levels of the anticancer drugs vinblastine and vincristine. The MIA pathway displays a complex architecture spanning different subcellular and cell-type localizations and is under complex regulation. As a result, the development of strategies to increase the levels of the anticancer MIAs has remained elusive. The pathway involves mesophyll specialised idioblasts where the late unsolved biosynthetic steps are thought to occur. Here, protoplasts of *C. roseus* leaf idioblasts were isolated by fluorescence-activated cell sorting, and their differential alkaloid and transcriptomic profiles were characterised. This involved the assembly of an improved *C. roseus* transcriptome from short- and long-read data, IDIO+. It was observed that *C. roseus* mesophyll idioblasts possess a distinctive transcriptomic profile associated with protection against biotic and abiotic stresses, and indicative that this cell type is a carbon sink, in contrast with surrounding mesophyll cells. Moreover, it is shown that idioblasts are a hotspot of alkaloid accumulation, suggesting that their transcriptome may hold the keys to the in-depth understanding of the MIA pathway and the success of strategies leading to higher levels of the anticancer drugs.

**Highlight:** Catharanthus mesophyll idioblasts are a hotspot of anticancer alkaloid accumulation. The idioblast transcriptome reveals commitment with stress responses and provides a roadmap towards the increase of anticancer alkaloid levels.

## Introduction

Understanding plant specialised metabolism is central to decipher plant defence strategies, as well as to fully exploit its diverse human applications, often hindered by metabolite low levels and limited availability of plant material. The advent of high-throughput sequencing has provided a way to tackle these hurdles, by enabling the thorough transcriptomic profiling of conditions, organs, tissues, or cell types associated with specific metabolic profiles, thus generating unprecedented insights into the genetic determinants of the architecture and dynamics of specialised metabolism (Dugé de Bernonville *et al.*, 2020; Courdavault *et al.*, 2021).

*Catharanthus roseus* (L.) G. Don (*Apocynaceae*) is the source of the low-level anticancer drugs vinblastine (VLB) and vincristine (VCR), and of the antihypertensive drug ajmalicine (Fig. 1). These drugs are monoterpenoid indole alkaloids (MIAs) and, despite intense research, the MIA pathway still hides several blindspots and remains defiant concerning VLB and VCR low levels. The biosynthesis of the dimeric VLB from the basic precursors tryptophan (indole moiety precursor) and geranyl pyrophosphate (monoterpenoid moiety precursor) involves nearly 30 enzymatic steps (Fig. 1). The pathway has been elucidated up to the coupling of the VLB monomeric building blocks, vindoline and catharanthine, into α-3’,4’-anhydrovinblastine (AVLB) (Costa *et al.*, 2008; Dugé de Bernonville 2015a; Kulagina *et al.*, 2022). However, the conversion of AVLB into the extremely low-level VLB and VCR remains elusive.

**Fig. 1.**
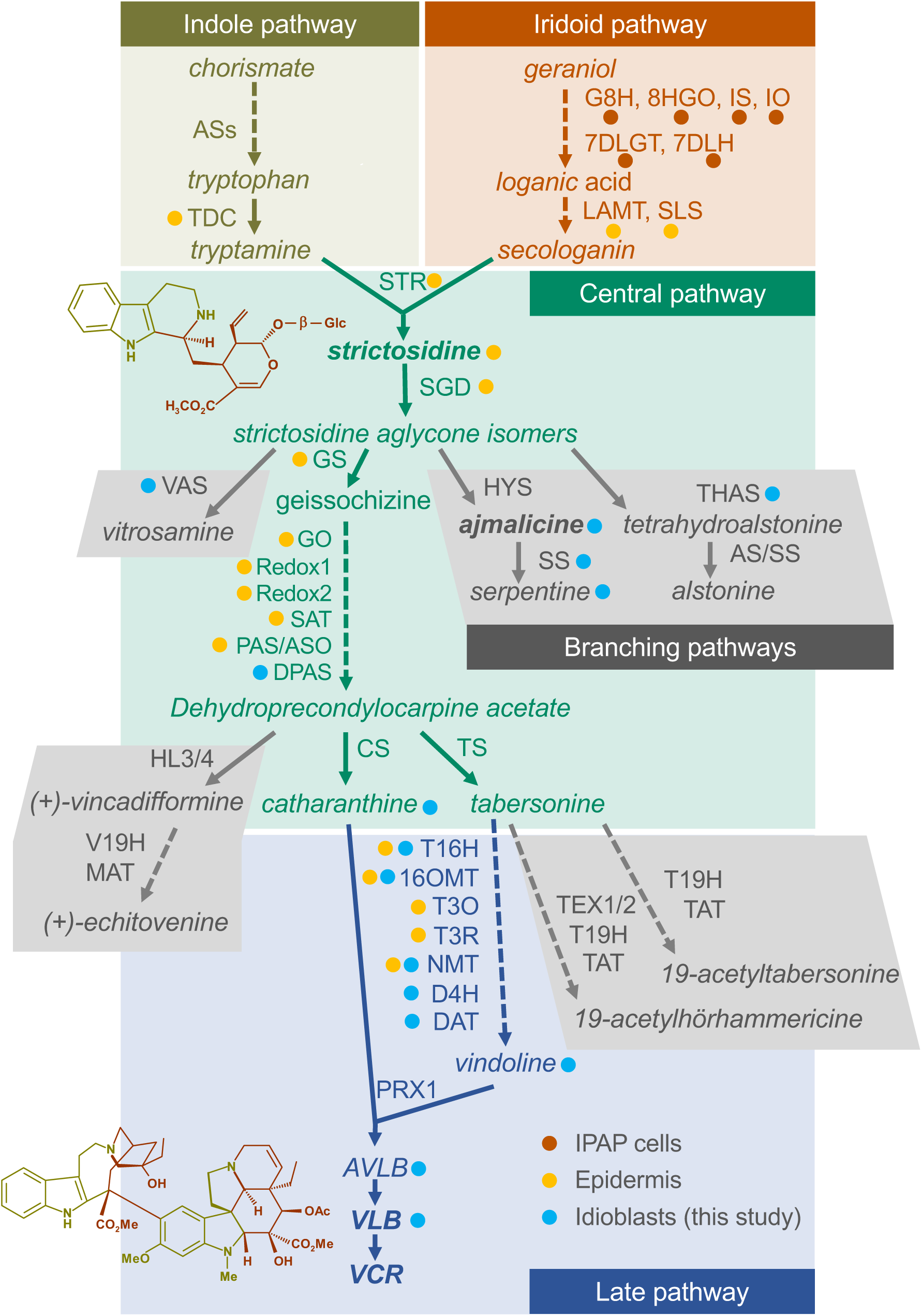
The MIA pathway in *C. roseus.* The indole and iridioid monoterpenoid pathways generate the indole precursor tryptamine and the monoterpenoid precursor secologanin, which condensate to yield strictosidine, the first MIA and the precursor of all MIA scaffolds. The central MIA pathway generates catharanthine and tabsersonine and is thought to occur in the epidermis. The late MIA pathway involves the conversion of tabersonine into vindoline, and the steps leading to the anticancer alkaloids VLB and VCR. Idioblast localization is represented by blue dots and is based on the results of this study. Epidermal and IPAP (internal phloem associated parenchyma) localization is represented by yellow and red-orange dots respectively, and has been previously published, corresponding to transcript localization by *in situ* analysis or presence in an epidermal RNA library, complemented in a few cases with protein localization using antibodies (St. Pierre *et al.*, 1999; Irmler *et al.*, 2000; Burlat *et al.*, 2004; Murata *et al.*, 2008; Guirimand *et al.*, 2011a; Guirimand *et al.*, 2011b; Geu-Flores *et al.*, 2012; Miettinen *et al.*, 2014; Qu *et al.*, 2015; Qu *et al.*, 2018a; Qu *et al.*, 2018b; Qu *et al.*, 2019). Structural formulae correspond to strictosidine (above) and VLB (below), with colours representing the pathway origin of each part of the backbone. The iridoid pathway: G8H, geraniol 8-hydroxylase; 8HGO, 8-hydroxygeraniol oxidoreductase; IS, iridoid synthase; IO, iridoid oxidase; 7DLGT, 7-deoxyloganetic acid glucosyltransferase; 7DLH, 7-deoxyloganic acid hydroxylase; LAMT, loganic acid O-methyltransferase; SLS, secologanin synthase. The indole pathway: AS, anthranilate synthase; TDC, tryptophan decarboxylase. The central pathway: STR, strictosidine synthase; SGD, strictosidine-β-D-glucosidase; GS, geissoschizine synthase; GO, geissoschizine oxidase; SAT, stemmadenine-O-acetyltransferase; PAS, precondylocarpine acetate synthase; DPAS, dihydroprecondylocarpine synthase; CS, catharanthine synthase; TS, tabersonine synthase. The late pathway: T16H, tabersonine 16-hydroxylase; 16OMT, tabersonine 16-O-methyltransferase; T3O - tabersonine 3-oxygenase; T3R - tabersonine 3-reductase; NMT, 16-methoxy-2,3-dihydro-3-hydroxytabersonine N-methyltransferase; DAT, deacetylvindoline-4-O-acetyltransferase; D4H, desacetoxyvindoline-4-hydroxylase; PRX1, class III peroxidase 1. Branching pathways: VAS, vitrosamine synthase; HYS, heteroyohimbine synthase; THAS, tetrahydroalstonine synthase; AS, alstonine synthase; SS, serpentine synthase; HL, hydrolase; V19H, vincadifformine-19-hydroxylase; MAT, minovincinine-19-O-acetyltransferase; TEX, tabersonine epoxidase; T19H, tabersonine 19-hydroxylase; TAT, 19-hydroxytabersonine 19-O-acetyltransferase.

MIA biosynthesis and accumulation in *C. roseus* is modulated by jasmonates and is under multilayer transcriptional regulation that has been partially deciphered, including the characterization of several transcription factors inducing and repressing the early pathways (Patra *et al.*, 2018, Schweizer *et al.*, 2018, Colinas *et al.*, 2021). Promising results were obtained recently with the characterization of the transcription factors ethylene responsive factor 5 (ERF5) and GATA binding protein 1 (GATA1), which, for the first time, enabled up-regulation of the late MIA pathway genes and alkaloids (Liu *et al.*, 2019; Pan *et al.*, 2019). Nevertheless, the transcriptional regulatory network of MIA biosynthesis is far from being fully understood.

The MIA pathway displays a surprisingly complex spatial architecture, spanning five subcellular compartments throughout four different cell types in *C. roseus* leaves, the organ where the dimeric anticancer VLB and VCR are specifically produced and accumulated (Courdavault *et al.*, 2014; De Luca *et al.*, 2014; Dugé de Bernonville 2015a; Kulagina *et al.*, 2022). The monoterpenoid moiety pathway (Fig. 1) starts in the chloroplasts and cytosol of internal phloem associated parenchyma (IPAP) leaf cells, after which it proceeds to leaf epidermis, where the monoterpenoid and indole final precursors, secologanin and tryptamine, are generated in the cytosol (Burlat *et al.*, 2004; Miettinen *et al.*, 2014). Secologanin and tryptamine are condensed inside the vacuole of epidermal cells by strictosidine synthase (STR) to yield strictosidine, the first MIA of the pathway and the central precursor of all MIA scaffolds (Fig. 1) (Guirimand *et al.*, 2011a). Many of the subsequent reactions appear to proceed in the epidermis, a localization that has also been observed for other alkaloid pathways and is consistent with the assumed defence role of these compounds (Guirimand *et al.*, 2011b; Qu *et al.*, 2019). The final steps of vindoline biosynthesis were shown to be specifically expressed in leaf laticifers and in specialized mesophyll cells – the idioblasts, both of which have also been shown to constitute leaf alkaloid accumulation targets (Yoder and Mahlberg, 1976; Mersey and Cutler, 1986; St. Pierre *et al.*, 1999; Carqueijeiro *et al.*, 2016; Yamamoto *et al.*, 2019). All multiple intra- and inter-cellular transmembrane transport events involved in the MIA pathway’s architecture are putative rate-limiting steps of MIA metabolic fluxes. However, so far, only a few of those events have been characterised (Carqueijeiro *et al.*, 2013; Yu and De Luca 2013; Payne *et al.*, 2017; Larsen *et al.*, 2017).

Leaf idioblasts, whose vacuoles are likely a final alkaloid accumulation target, are potentially the home of important VLB/VCR biosynthetic steps and of transmembrane transport events with a high impact in the metabolic flux of the MIA pathway. In this study, *C. roseus* leaf idioblast protoplasts were isolated using a fluorescence-activated cell sorting (FACS) methodology (Carqueijeiro *et al.*, 2016), and the characterization of the differential alkaloid and transcriptomic profile of this cell type was performed. The sequence data obtained enabled the generation of a comprehensive transcriptome, IDIO+, which represents a new valuable tool for the species. It is shown that the *C. roseus* mesophyll idioblasts possess a unique transcriptomic profile, deeply implicated in protection against biotic and abiotic stresses, and displaying low expression of genes involved in photosynthesis, in comparison with common mesophyll cells. It is also shown that idioblasts accumulate much higher levels of MIAs than other leaf cells, suggesting that their transcriptome information may be decisive to the understanding of the VLB/VCR pathway towards manipulation of *C. roseus* for higher levels of those anticancer drugs.

## Materials and methods

### Plant material

*C. roseus* (L.) G. Don ‘Little Bright Eye’ plants were grown at 25 °C in a growth chamber under a 16 h / 8 h photoperiod, using white fluorescent light with a photon medium intensity of 70 μmol m^-2^ s^-1^. Seeds were acquired from B&T World Seeds.

For the indoor / outdoor experiment, one group of *C. roseus* mature flowering plants were kept in the growth chamber, while an equivalent group was transferred outdoors, under natural environmental conditions and exposed to direct sunlight during ∼ 6 months. Leaves of the first, second, third and fourth pair (counting from the shoot apical meristem) of three plants from each set were collected around 14.00h, in the beginning of September, in a hot (above 30°C) sunny day. After the removal of central veins, leaves were macerated with liquid nitrogen and part was stored at -80°C for RNA extraction, while the remaining was freeze-dried and stored at -20°C for alkaloid extraction.

### Isolation of leaf protoplasts and FACS

Six-month-old plants were used for protoplast isolation and three biological replicates were performed. Each replicate contained 1.5 to 2 g of fully developed healthy young leaves from the second to the fourth pair (counting from the shoot apex), collected from three mutually exclusive groups of three to five plants. Protoplast isolation was performed as previously described in Carqueijeiro *et al.* (2013). The FACS methodology is described in Carqueijeiro *et al.* (2016) and Guedes *et al.* (2022). Matching leaf samples were also collected and processed as described in the plant material section.

### Alkaloid extraction and analysis by HPLC-Mass Spectrometry

Fresh material was freeze-dried immediately after being collected. Leaf samples contained 70 to 180 mg FW. The number of cells used for each sorted sample was 200,000 for total protoplasts, 300,000 for mesophyll protoplasts and 30,000 for idioblast protoplasts. Alkaloids were extracted by adding 150 or 500 µL of HPLC grade methanol to cell or leaf samples respectively. Papaverine was added at this stage as an internal control: 17,5 or 35 µg to cell or leaf samples respectively, using a 2,5 mg mL^-1^ methanolic solution. Samples were resuspended and extracted by vortexing during 2 min, followed by sonication for 30 min, and centrifugation at 10,000 *g* for 10 min. Supernatants were filtered and analysed by HPLC-Electrospray Ionization-Tandem Mass Spectrometry as described in Carqueijeiro *et al.* (2016). All alkaloid standards were obtained from Sigma Aldrich, except AVLB, which was obtained from Pierre Fabre. Retention times (RT) and masses were as follows: serpentine, RT = 8.03 min, *m/z* 349,15; papaverine, RT = 10.98 min, *m/z* 340,15; vindoline, RT = 19,30 min, *m/z* 457,23; vincristine, RT = 26.12 min, *m/z* 825,41; catharanthine, RT = 29,00 min, *m/z* 337,19; ajmalicine, RT = 34.04 min, *m/z* 353,19; vinblastine, RT = 38,63 min *m/z* 811,43; anhydrovinblastine, RT = 49,20 min, *m/z* 793,42.

### RNA extraction, library preparation and sequencing

RNA was extracted from paired samples of whole leaves, total protoplasts (200,000 cells), sorted idioblast protoplasts (50,000 cells) and sorted mesophyll protoplasts (120,000 cells). To purify RNA from the three protoplast populations, a solution of 1% (v/v) ß-Mercaptoethanol in RNeasy Lysis Buffer (Quiagen) was added in a volume ratio of 3.5:1 of cell suspension. Cells were disrupted by vigorous vortexing, flash-frozen in liquid nitrogen and stored at -80°C. RNA extraction was performed with the RNeasy Micro Kit (Qiagen) protocol in agreement with manufacturer’s instructions. For whole leaf samples, processed as described in the plant material section, RNA was extracted with the RNeasy Plant Mini Kit (Qiagen) following manufacturer’s instructions. All RNA extracts were precipitated with glycogen RNA-grade (Thermo Scientific), according to the manufacturer’s protocol. Due to RNA degradation, one leaf sample was discarded. For the FACS experiment, libraries were prepared using the Smart-seq2 kit following the manufacturer’s protocol (Picelli *et al.*, 2013). For the indoor / outdoor experiment, libraries were prepared using the TruSeq RNA Library Prep Kit. Sequencing was performed with Illumina HiSeq1500 on pair-end mode (2×125 bp) for 250 cycles. Long read libraries were prepared with Nanopore SQK-PCB109 kit following the manufacturer’s protocol. Sequencing was performed on single-end mode with a MinION sequencer using an R9.4.1 flow cell for 71 hours.

### Data quality control and filtering

Illumina raw data was processed with Trimmomatic (version 0.39) (parameters: ILLUMINACLIP:2:30:10 LEADING:3 TRAILING:10 SLIDINGWINDOW:4:15 MINLEN:50) to remove adaptors and low-quality bases, and to filter reads with ≤ 50 base pairs (Bolger *et al.*, 2014). For Nanopore data, basecalling and demultiplexing was performed using the Guppy v4.0.14 (Oxford Nanopore Technologies—ONT, Oxford, UK) high accuracy model. All reads were oriented based on barcode sequences using Pychopper (version 2.4, https://github.com/Nanoporetech/pychopper). During this process, barcode sequences were trimmed, low-quality reads were removed, and reads with < 50 bp were discarded. Both “oriented” and “rescued” reads were used for downstream analysis. Data quality and statistics was verified using the FastQC tool (version 0.19.11, https://www.bioinformatics.babraham.ac.uk/projects/fastqc/).

### Genome-guided transcriptome assembly and annotation

Following read filtering, all samples were mapped to the *C. roseus* genome v2 (Franke *et al.*, 2019; https://datadryad.org/stash/dataset/doi:10.5061/dryad.08vv50n). For Illumina data, reads were aligned using HISAT2 (version 2.2.0) using the “-dta” option enabled (Kim *et al.*, 2019). Following this step, a transcriptome for each sample was assembled using Stringtie (version 2.1.4) setting the minimum isoform abundance *per locus* to 0.05 (Kovaka *et al.*, 2019). Nanopore long reads were mapped to the *C. roseus* genome v2 using minimap2 (version 2.15-r905 using parameters -ax splice -uf; Li, 2018), and the primary alignments were used for assembly with Stringtie (using parameters -rf -L).

The Mikado pipeline (version 2.1.0) was then used to merge Stringtie runs and remove redundant sequences (Venturini *et al.*, 2018). Several metrics related to each gene model were computed to aid in improving the final transcriptome reconstruction. Sequences were subjected to open reading frame calling using TransDecoder (version 5.5.0; Haas *et al.*, 2013) and to homology searches against the Swiss-Prot viridiplantea database using DIAMOND (version 2.0.6 using parameters -e 0.00001 -sensitive -max-target-seqs 50; Buchfink *et al.*, 2015). High quality splicing junctions were obtained from the Illumina alignments using Portcullis (Mapleson *et al.*, 2018). The CDF97 transcriptome (Dugé de Bernonville *et al.*, 2015b), *C. roseus* complete genes downloaded from NCBI (accessed on 14-02-2021) were mapped to the genome v2 reference annotation using minimap2, and sequences longer than 2,500 base pairs containing two or more hits (>90% query coverage and >80% identity) were identified as putative chimeric reference gene models. The final set of transcripts was obtained using Mikado pick with the “plant.yaml” scoring file, with the above mentioned fusion gene models being penalised. The genome browser Jbrowser (Buels *et al.*, 2016) was used for visualisation of resulting gene-models and manual curation was performed when necessary. GTF and FASTA files including all the mRNA gene models were generated (Supplementary Datasets S1, S2). Gffcompare (Pertea and Pertea, 2020) was used for identification of novel genes, by comparing the final transcriptome with the reference gene models (Supplementary Table S1). BUSCO (Benchmarking set of Universal Single-Copy Orthologs; version 5.0.0) was used to assess the completeness of the final transcriptome using the eudicots_db10 dataset as queries. The full IDIO+ transcriptome (TSA, transcriptome shotgun assembly), including coding, non-coding and splicing variants has been deposited at DDBJ/EMBL/GenBank under the accession GKIJ00000000. The version described in this paper is the first version, GKIJ0100000.

The functional annotation of coding protein sequences was performed using both the pannzer2 (Toronen *et al.*, 2018) web server (http://ekhidna2.biocenter.helsinki.fi/sanspanz/) and the InterproScan software (version 5.50-84.0) (Quevillon *et al.*, 2005) with default parameters (Supplementary Dataset S3). Only the representative transcript isoform for each gene (defined as the higher scoring Mikado transcript for each gene) was annotated. Gene ontology (GO) terms were obtained from pannzer2 annotation. Enzymes, transporters, and transcription factors from the MIA pathway were annotated manually, with the GO annotation being retrieved from the Uniprot database (Liu *et al.*, 2017).

### Quantification and exploratory analysis

Following mapping, quantification of short reads was obtained with featureCounts (version 2.0.1, parameters: -p -t exon -g gene_id) using the mRNA gene models gtf file (Supplementary Dataset S1). Transcripts per million (TPM) values were calculated using the formula: 10^6^ x (reads_mapped_to_transcript / transcript_length) / Sum (reads_mapped_to_transcript / transcript_length), where reads_mapped_to_transcript is the total number of reads mapped to a gene and transcript length is the size of the gene as outputted by featureCounts. Principal component analysis (PCA) and hierarchical clustering based on sample correlation was performed on short read data. Count data was imported to R, normalized using the *DESeq2* package (version 1.28.1) and transformed using the *rlog* function (Love *et al.*, 2014). For the PCA, only the top 500 genes, in terms of variance, were considered. Spearman correlation was used for the hierarchical clustering and the figure was generated using the *pheatmap* package. The RNA-seq data have been deposited in NCBI’s Gene Expression Omnibus (Edgar *et al.*, 2002) and are accessible through GEO Series accession number GSE217852 (https://www.ncbi.nlm.nih.gov/geo/query/acc.cgi?acc=GSE217852).

### Differential gene expression analysis

Differential gene expression (DGE) analysis was performed using the *DESeq2* package (Soneson *et al.*, 2015). For the analysis, idioblast, mesophyll and total protoplasts samples, as well as indoor and outdoor plant samples from leaf pairs 1 and 4 (short reads) were incorporated into the statistical model. Leaf samples from the FACS experiment were excluded from the model due to a higher within-group variance (Supplementary Fig. S1). Only genes with adjusted p-values ≤ 0.05 and log2 fold change ≥ 1 were considered differentially expressed. The results of DGE are included in Supplementary Tables S2-S4. The TPM expression levels for all the samples used can be found in Supplementary Dataset S4.

### Weighted gene co-expression network analysis

Expression levels from mesophyll and idioblast protoplast samples as well as from leaf pair 1 and 4 from plants growing indoors / outdoors were used to build a co-expression network using the R package *WGCNA* (version 1.69; Langfelder and Horvath, 2008). To filter genes with low expression levels, count data was first normalized into count per million (CPM). Only genes with > 1.5 CPM in at least six of the samples were considered for network construction. The filtered gene matrix with raw counts was then subjected to normalization using *DESeq2* (Soneson *et al.*, 2015) and was subjected to variance-stabilizing transformation (Anders and Huber, 2010). For the signed network construction, an adjacency matrix was computed by raising to a power of 20 (soft threshold) the Pearson correlation between the expression profiles of every pair of genes (Supplementary Fig. S2). A topological overlap matrix was then computed based on the adjacency values and the network was constructed using the corresponding dissimilarity matrix. Gene modules were identified using the *cutreeDynamic* function, with the minimum module size set to 30. Resulting modules were clustered using a representative expression profile (the module “eigengene”), and correlated modules with a maximum height of 0.15 were merged. Enrichment analysis for each module was performed as described below using the network genes as the background annotation.

For module-trait correlation analysis, the measurements of catharanthine, vindoline, AVLB and VLB in idioblast and common mesophyll protoplast samples were converted from ng/cell into ng/mg of dry weight. It was assumed that the alkaloid content of total protoplasts and leaves was roughly the same, and the average ratio between pair-wise leaf/protoplast samples was applied to idioblast and common mesophyll protoplast samples to estimate values in ng/mg dry weight. Module-trait correlation analysis was performed by computing the Pearson correlation between module eigengenes and alkaloid levels. Correlations with a false discovery rate (FDR) adjusted p-value < 0.05 were considered significant (Benjamini and Hochberg, 1995).

### Enrichment Analysis

Enrichment analysis in GO terms was performed using the *topGO* package version 2.40 (Alexa *et al.*, 2006) for both upregulated and downregulated gene sets. The background annotation was defined as the full list of genes expressed in either idioblast or common mesophyll protoplasts. For the analysis, GO terms with less than 5 annotated genes were excluded, to prevent statistical artefacts. The “weigth01” algorithm was used during the analysis. The Fisher exact test was used to calculate significance for all tests, with p-values ≤ 0.01 being considered as statistically significant.

### Phylogenetic analysis

For phylogenetic analysis, the dicots PLAZA portal (version 4.5; https://bioinformatics.psb.ugent.be/plaza/versions/plaza_v4_5_dicots/) was surveyed to obtain all protein sequences for each studied gene family, from the genomes of *Arabidopsis thaliana*, *Amborella trichopoda* and *Oryza sativa* (Van Bel *et al.*, 2012). In addition, protein sequences of members with known functions in specialized metabolism, as concluded from a literature survey, were retrieved from GenBank. Protein sequence data was aligned with the *C. roseus* candidate sequences using MAFFT (version 7.475, default parameters, Katoh and Standley, 2013) and the most fitting evolutionary model was determined with SMS version 1.8.4 (Lefort *et al.*, 2017) using BIC (Bayesian information criterion) as the selection criterion (Supplementary Table S5). Phylogenetic analyses were performed on the CIPRES portal (http://www.phylo.org/) (Miller *et al.*, 2011) using RaxML version 8.2.12 (Stamatakis, 2014), with 1000 rapid bootstraps (BS). Resulting trees were visualized and edited using FigTree (version 1.4.4; http://tree.bio.ed.ac.uk/software/figtree/).

## Results

### Anticancer alkaloids are specifically accumulated in idioblasts

To establish the precise partaking of *C. roseus* leaf idioblasts in the MIA pathway, a highly enriched population of leaf idioblast protoplasts was generated by FACS (Fig. 2A), using a protocol previously developed in our lab (Carqueijeiro *et al.*, 2016). The procedure also enabled the generation of pure populations of common mesophyll cell protoplasts from the same cell samples.

**Fig. 2.**
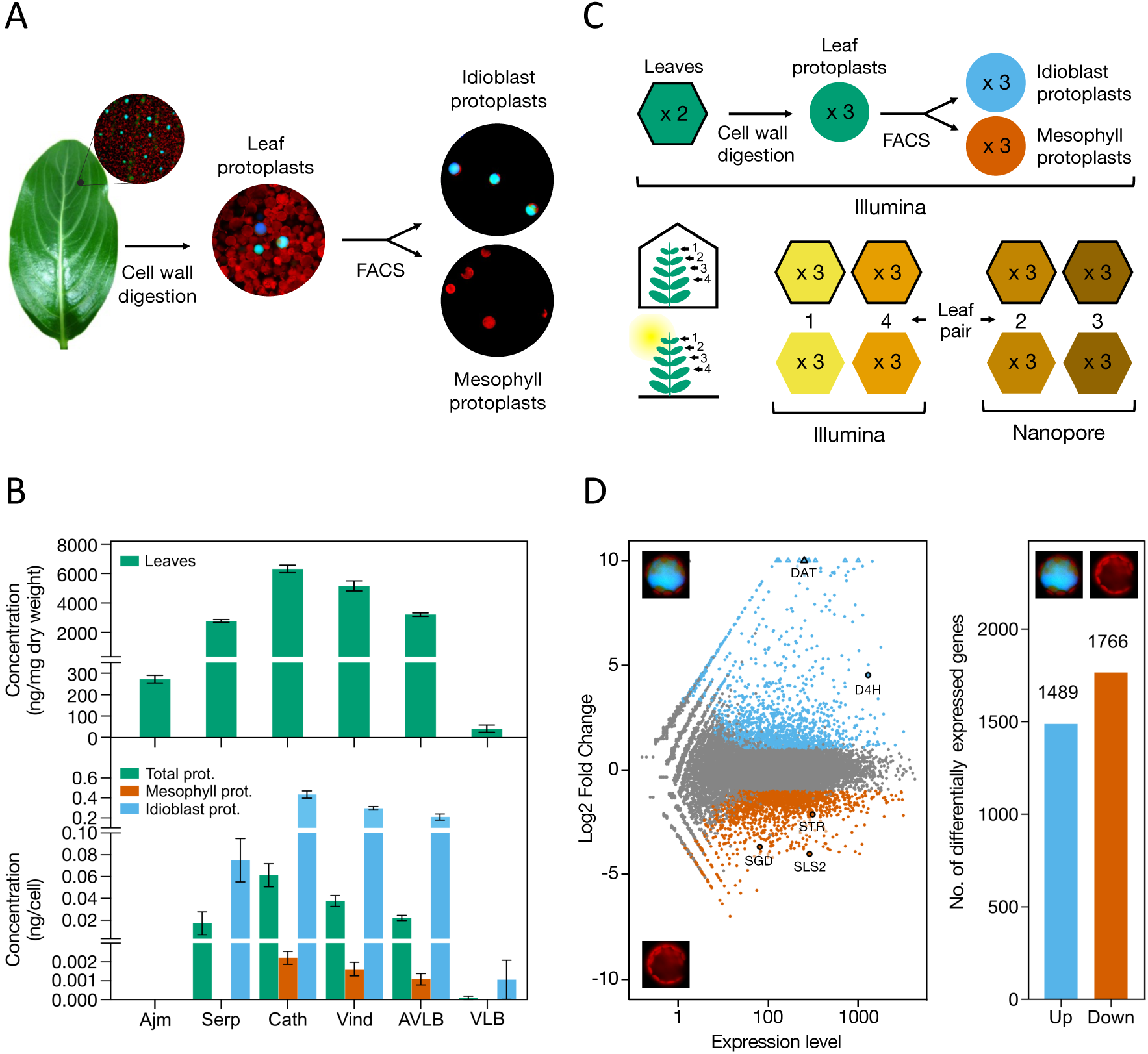
(A) Outline of the experimental setup used to generate populations of *C. roseus* leaf idioblast cells and common mesophyll cells. (B) Levels of the main alkaloids detected in leaves, total protoplasts, sorted common mesophyll protoplasts and sorted idioblast protoplasts. N = 3. (C) Experimental design used to generate RNA-seq data for transcriptome assembly. Samples/replicates from the idioblast experimental setup displayed in A were used for Illumina sequencing. Samples/replicates from leaf pairs 1 and 4 (counting from the shoot apex) of an experiment including one group of control plants grown in a growth chamber (indoor plants) and another group of plants grown outdoors whilst submitted to natural light and temperature conditions (outdoor plants), were used for Illumina sequencing. Samples/replicates from leaf pairs 2 and 3 of the latter experiment were used for Nanopore sequencing. Hexagons represent leaf samples and circles represent protoplast samples. (D) Summary plots of differentially expressed genes in idioblast protoplasts compared with protoplasts of common mesophyll cells. Left: *MA*-plot showing idioblast upregulated genes in blue, idioblast downregulated genes in red-orange, and non-differentially expressed genes in grey. Right: bar plot showing the numbers of differentially expressed genes. Genes with absolute log_2_ fold change > 1 and adjusted p-values < 0.05 were considered as differentially expressed.

The protoplasts of idioblasts and common mesophyll cells obtained with the applied methodology have been previously shown to retain their *in planta* distinctive identities (Carqueijeiro *et al.*, 2016), and may therefore be used to investigate the precise metabolic and transcriptomic profiles of these two leaf cell types. Idioblast populations contained approx. 90% of cells depicting the intense blue/green autofluorescence and high refraction of the vacuolar content characteristic of *C. roseus* leaf idioblasts, while mesophyll cell populations were constituted exclusively by this cell type. Fig. 2B shows the levels of the alkaloids detected in the leaves of the plants used and across the separate protoplast populations. The levels of all the alkaloids detected were more than hundredfold in idioblasts in comparison with common mesophyll cells - from 183 fold for vindoline to 196 fold for catharanthine.

### A novel transcriptome assembly using long-read data improves current gene models

The observed importance of idioblasts for alkaloid accumulation in *C. roseus* leaves (Fig. 2B) prompted the investigation of the differential transcriptomic profile of this cell type, aiming to identify candidate genes for key biosynthetic, transport and transcriptional regulatory events of the MIA pathway, as well as to build up a global picture of the biology of this cell type. Idioblasts are under-represented within leaf tissues and, therefore, their transcripts may not have been detected in previous *C. roseus* transcriptomic projects, which were the main sources for the identification of coding sequences in the *C. roseus* genomes v1 and v2 (Kellner *et al.*, 2015; Franke *et al.*, 2019). As such, a genome-guided transcriptome assembly was performed, using RNA-seq short-read data obtained from the same samples as the ones analysed for alkaloid levels, complemented with RNA-seq data generated from a second experiment (Fig. 2C). In this experiment, a group of control plants was grown in a growth chamber (indoor plants) while another group was grown outdoors, submitted to natural light and temperature conditions (outdoor plants). The dataset for this second experiment included Nanopore long-read sequencing of 12 independent biological samples, significantly enhancing the transcriptome assembly with putative full-length transcript information.

The RNA-seq experiment design illustrated in Fig. 2C generated a short-read dataset containing 432 million paired-end reads (2×125 bp) across 23 biological samples, and a long-read dataset containing 12.6 million reads across 12 biological samples with a median length of 479 bp and a N50 of 731 bp. Consistency in alignment ratio was observed across all samples, with over 90% of the short-read data and 84% of the Nanopore data mapping to the genome v2. An overview of the data can be found in Supplementary Table S6.

The approach used for the genome-guided transcriptome assembly is summarised in Supplementary Fig. S3. After the initial assembly and analysis, several erroneous gene-models were detected not only on the individual Stringtie runs but also on the original reference annotation (Supplementary Fig. S4). These artefacts included transcript fusion, fragmented transcripts and incorrect exon-intron chains. To minimise this issue, Mikado was employed to merge the independent sample assemblies and refine the resulting transcripts (Venturini *et al.*, 2018). This approach corrected several gene-models, including models present in the genome v2 reference annotation. The novel assembly contained 59,496 transcriptomic isoforms across 38,788 genes, of which 35,722 were predicted to have an open reading frame (Supplementary Table S7). Among these, 1,128 were novel gene loci previously undetected in the *C.roseus* genome v2 reference annotation (Supplementary Table S1). Most significantly, 567 of these new genes are expressed in idioblasts. This novel transcriptome assembly is hereafter named IDIO+.

Assessment with BUSCO detected 96% of the eudicots core orthologs as complete and 1.3% as fragmented in IDIO+, which represents a significant improvement over the annotation of the reference genome v2, whose scores are 90.6% complete and 3.5% fragmented (Supplementary Table S7; Franke *et al.*, 2019; Seppey *et al.*, 2019). The IDIO+ BUSCO scores are similar to the CDF97 consensus transcriptome reported by Dugé de Bernonville *et al.* (2015b).

Protein coding genes were annotated with panzer2 and resulted in GO terms being assigned to 34%, 35.1% and 35.5% of the genes for Biological Process, Molecular Function and Cellular Component, respectively (Toronen *et al.*, 2018). Functional domains and protein families were also annotated using Interpro (Blum *et al.*, 2021) and resulted in 59.9% of the genes being annotated in at least one of the member databases. The overall annotation rate across both methods was 61%. Since the MIA pathway is central to this study, annotation of this pathway’s enzymes, transporters and transcription factors was performed manually in IDIO+ (Supplementary Table S8).

### Differential gene expression reveals a clear specialization of idioblasts

The improved *C. roseus* reference gene-models of IDIO+ were used to perform the characterization of the transcriptomic landscape of idioblasts. Principal component analysis (PCA) and hierarchical clustering based on the gene expression profiles revealed that idioblasts and common mesophyll cells have clearly distinct transcriptomic profiles (Supplementary Figs S5, S6). This allowed the implementation of DGE analysis of idioblast protoplasts, using the mesophyll protoplast samples as a reference.

The idioblastome included 1,489 genes overexpressed in this cell type, relative to common mesophyll cells, while 1,776 genes were underexpressed (Fig. 2D). The full list of differentially expressed genes can be found in Supplementary Tables S2-S4. The top 25 idioblast upregulated genes ranked by log2 fold change and the top 25 genes ranked by expression levels are presented in Tables 1 and 2, respectively. The top upregulated / highly expressed idioblast genes include genes involved in the biosynthesis of MIAs (e.g.: deacetylvindoline-4-O-acetyltransferase, *DAT*; desacetoxyvindoline-4-hydroxylase, *D4H*; and tetrahydroalstonine synthase, *THAS*), as well as enzymes, transporters and a transcription factor belonging to gene families implicated in the MIA pathway. Other suggestive top idioblast genes include oxygenases, major latex proteins, dehydrins, patatin, amongst others, which are either related with specialised metabolism or stress.

To further understand the underlying processes and roles played by idioblasts, a GO term enrichment analysis was performed for both the up and downregulated genes (Fig. 3). For the GO category biological process, the top enriched term is “transmembrane transport” for both up and downregulated genes, indicating an important differential specialization of idioblasts. Upregulated “transmembrane transport” represents the term with the highest number of annotated genes (105) in the category biological process, which further includes enrichment in “xenobiotic transport”, “purine nucleoside transport” and “copper ion transmembrane transport”. Transporter activities are also the bulk of GO terms enriched in idioblasts for the category molecular function, with “integral component of membrane” being the top enriched term for the category cellular component, further indicating a strong distinctiveness of idioblast membranes.

**Fig. 3.**
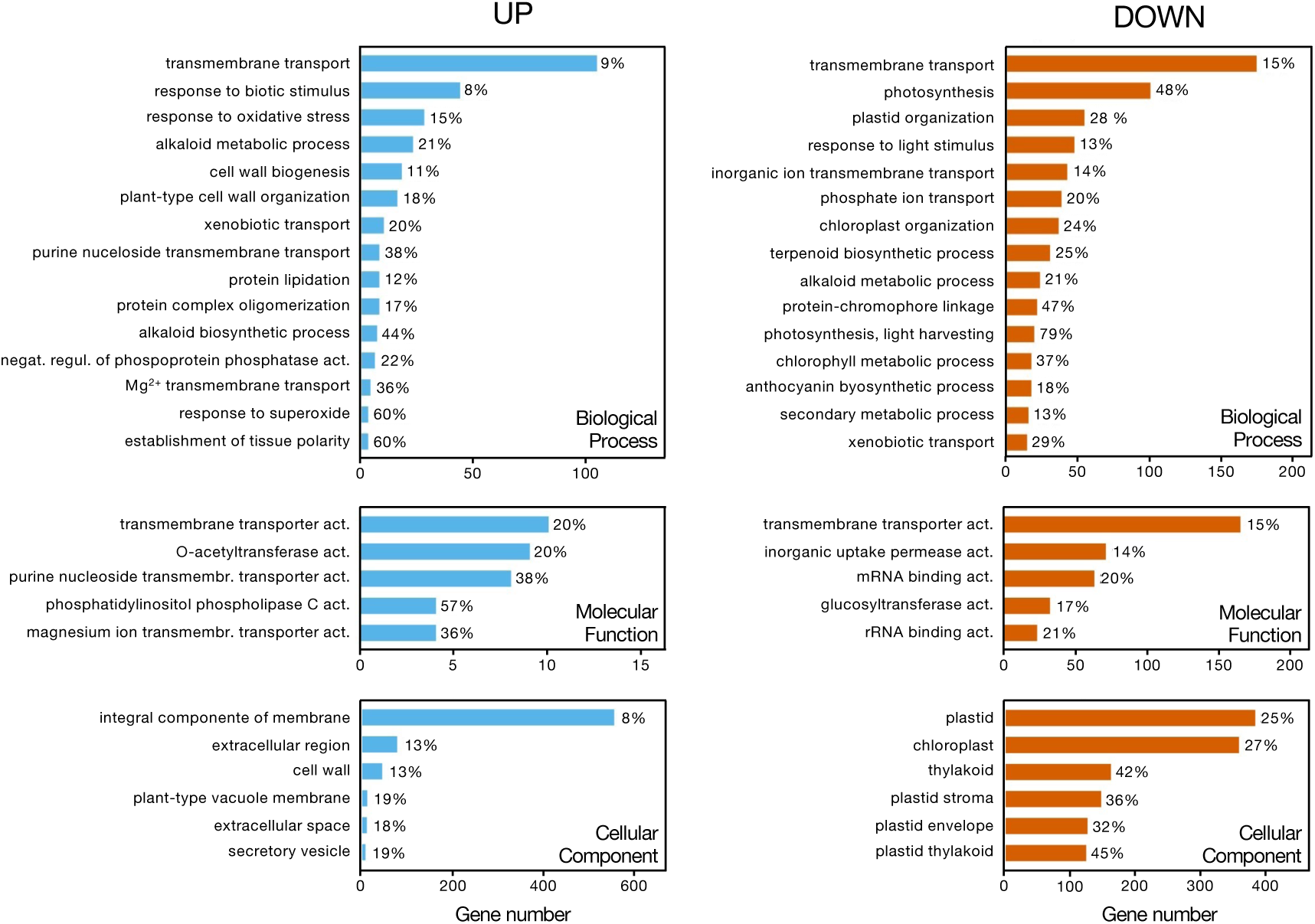
Enrichment analysis of GO terms associated with genes upregulated (left, blue) and downregulated (right, red orange) in idioblast protoplasts compared with common mesophyll cell protoplasts. The percentage of up or downregulated genes from each GO term, in relation to the background annotation (list of genes expressed in either idioblasts or common mesophyll protoplasts), is represented next to the respective bar. When a category included more than 15 significantly enriched terms, only the top 15 (ordered by the number of genes up or downregulated) were included.

Enrichment analysis of idioblast upregulated genes further revealed an overrepresentation of the term “responses to biotic stimulus”, which included genes putatively involved in “defence response to fungus” and “defence response to bacterium” (Fig. 3A). The related term “response to oxidative stress” is also enriched in idioblasts. The terms “alkaloid metabolic process” and “alkaloid biosynthetic process” appear overrepresented in idioblasts, although “alkaloid metabolic process” is also enriched in the downregulated genes. Another idioblast enriched term worth mentioning is “establishment of tissue polarity”, which potentially includes genes involved in the differentiation of this cell type. Several GO terms related with the cell wall are also enriched in idioblasts, including “cell wall biogenesis” and “cell wall thickening”.

The top enriched GO terms among idioblast downregulated genes are mostly related with photosynthesis and chloroplast processes / components (Fig. 3B).

### Gene co-expression modules show correlation with alkaloid traits

The RNA-seq data of this study originated from two experimental setups: i) the FACS isolation of idioblast protoplasts (Fig. 2A, C) and ii) the collection of different leaf pairs from plants growing in a growth chamber (indoor plants), and plants submitted to natural light and temperature conditions (outdoor plants) (Fig. 2C). Idioblast protoplasts presented radically higher alkaloid levels than their mesophyll counter parts, and the samples from the indoor/outdoor experiment also presented a significant change in their alkaloid profiles, with the leaves of outdoor plants showing a significant increase in the levels of the dimeric AVLB, VLB and VCR, associated with a decrease in the levels of the monomeric precursors vindoline and catharanthine (Supplementary Fig. S7). Therefore, the extended RNA-seq dataset of this study was used to generate co-expression modules using weighted gene co-expression network analysis, followed by module-trait analysis inputting the levels of the alkaloids vindoline, catharanthine, AVLB and VLB. PCA of gene expression levels of all the different samples showed that leaf data from the two experiments dissociated well from protoplast samples (Supplementary Fig. S1). After network construction, module detection and clustering, 12 gene modules were obtained, and were then used for module-trait analysis (Fig. 4A and Supplementary Table S9). Overall, results show that catharanthine, vindoline, and AVLB levels present a very similar correlation pattern with the different gene modules, (Fig. 4A). In contrast, VLB levels show a strikingly different pattern of correlation with the gene modules, mostly anti-correlated with catharanthine, vindoline and AVLB patterns.

**Fig. 4.**
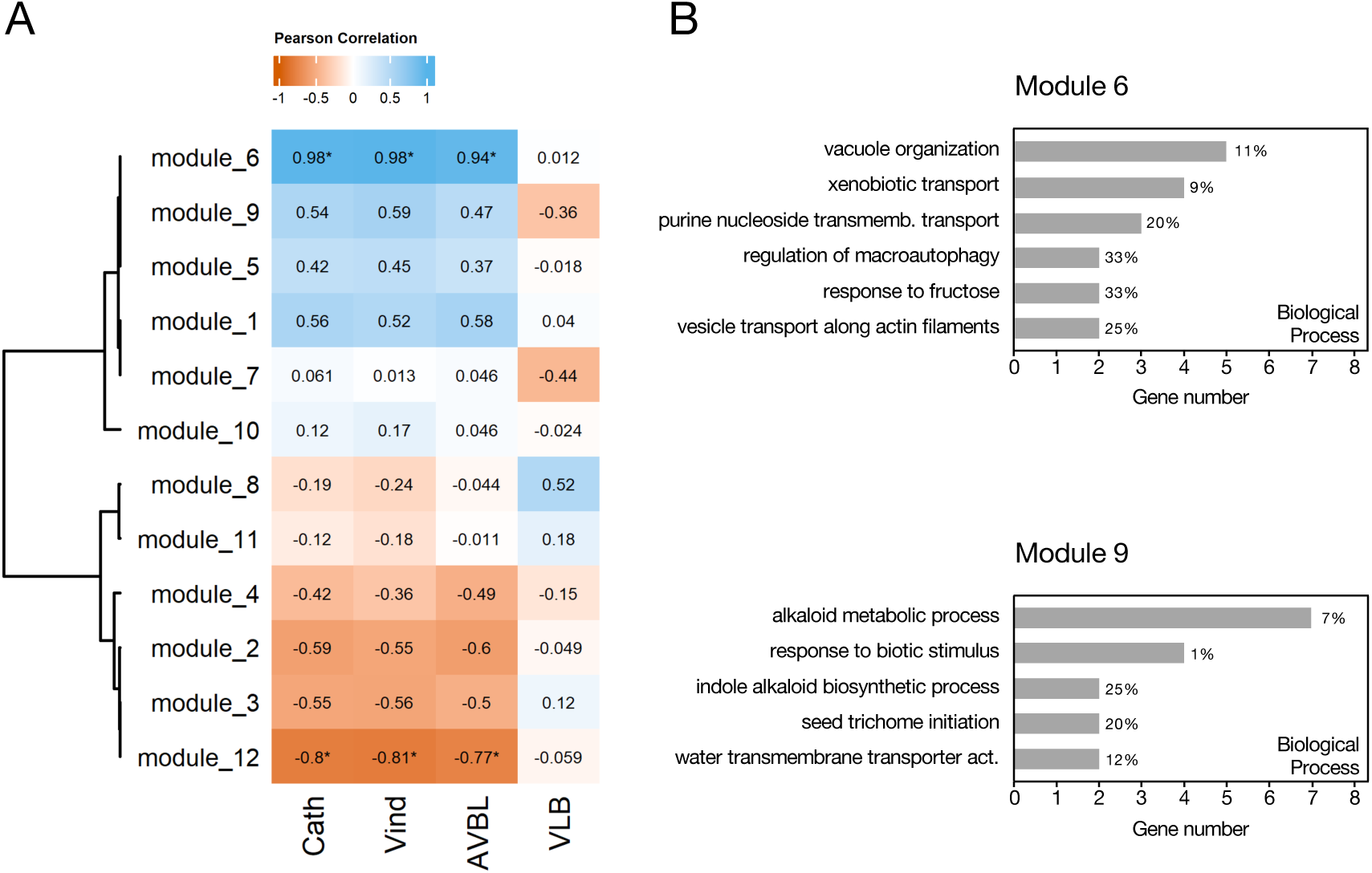
(A) Module-trait correlation analysis of gene co-expression modules with vindoline, catharanthine, AVLB and VLB levels. The clustered heatmap represents correlations between module eigengenes and alkaloid levels, with blue representing positive correlations and red orange representing negative correlations. Statistically significant correlations are marked with * (adjusted p-value < 0.05). (B) Enrichment analysis of GO terms (Biological Process) of module 6 and module 9 genes. The percentage of genes from each GO term present in the module, in relation to the background annotation (list of genes included in the co-expression analysis), is represented next to the respective bar.

Over 70% of the genes in modules 6 and 9 are upregulated in idioblasts (Supplementary Table S9), implying that they might be related to processes specific to these specialised cells. Module 6 presents a significant positive correlation with catharanthine (0.98), vindoline (0.98) and AVLB (0.94) traits, suggesting a key role in the regulation of MIA levels (Fig. 4A). Despite the high correlation, dihydroprecondylocarpine synthase (*DPAS*) was the only MIA gene found in this module. Module 6 has an overrepresentation of genes related to “xenobiotic transport” and “purine nucleoside transport” (Fig. 4B), corresponding, respectively, to multidrug and toxic compound extrusion (MATE) and purine permease (PUP) family transporters, which have been implicated in alkaloid transmembrane transport and may be determinants of MIA metabolic flux. Module 9 is closely correlated with module 6 and includes 3 vindoline pathway genes (tabersonine 16-O-methyltransferase - *16OMT*, *DAT* and *D4H*). However, the module-trait correlation analysis revealed that this module is not significantly correlated with alkaloid levels (Fig. 4A). GO term analysis revealed that module 9 is enriched in genes involved in alkaloid biosynthesis and response to biotic stimulus (Fig. 4B).

### Differential gene expression confirms specialization of idioblasts in the late MIA pathway

The results obtained for the differential expression in idioblasts of genes involved in the biosynthesis of the anticancer VLB are shown in Fig. 5. As already shown in Table 1 above, *D4H* and *DAT* (Fig. 1), responsible for the two last steps of vindoline biosynthesis, are strongly upregulated in the idioblast cell populations generated in this study. Five of the seven enzymatic steps involved in the late vindoline biosynthetic pathway are upregulated in idioblasts: tabersonine 16-hydroxylase 2 (*T16H2*), tabersonine 16-O-methyltransferase (*16OMT*), 3-hydroxy-16-methoxy-2,3-dihydrotabersonine N-methyltransferase (*NMT*)*, D4H* and *DAT*, responsible for the first, second, fifth, sixth and seventh steps respectively (Fig. 1). The fourth step, tabersonine 3-reductase (*T3R*), is downregulated, and no expression levels were detected for the third step, tabersonine 3-oxygenase (*T3O*), in mesophyll or idioblast protoplasts. Class III peroxidase 1 (*PRX1*), thought to be responsible for the dimerization reaction yielding AVLB (Costa *et al.*, 2008), is downregulated in idioblasts. All other parts of the MIA pathway leading to VLB are either downregulated in idioblasts or not differentially expressed, except *DPAS*, which is upregulated.

**Fig. 5.**
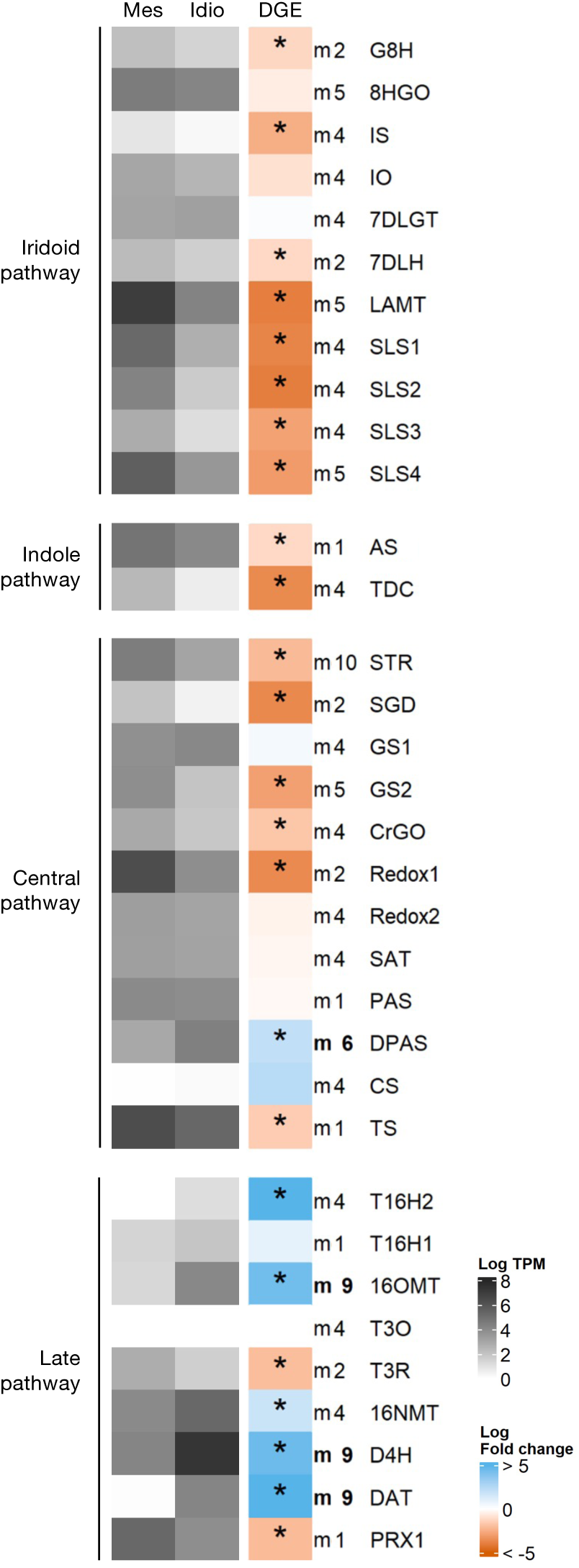
Expression analysis of genes involved in the biosynthesis of the anticancer VLB. Heatmap grey colours represent the mean log_2_ TPM of three independent biological samples. Log_2_ fold change for each gene is represented next to the expression level, with blue and red orange representing respectively overexpression and underexpression in idioblast protoplasts compared with mesophyll protoplasts. Significant differential expression is marked with a *. The iridoid pathway: *G8H*, geraniol 8-hydroxylase; *8HGO*, 8-hydroxygeraniol oxidoreductase; *IS*, iridoid synthase; *IO*, iridoid oxidase; *7DLGT*, 7-deoxyloganetic acid glucosyltransferase; *7DLH*, 7-deoxyloganic acid hydroxylase; *LAMT*, loganic acid O-methyltransferase; *SLS1* - secologanin synthase 1, *SLS2* - secologanin synthase 2, *SLS3* - secologanin synthase 3, *SLS4* - secologanin synthase 4. The indole pathway: *AS*, anthranilate synthase; *TDC*, tryptophan decarboxylase. The central pathway: *STR*, strictosidine synthase; *SGD*, strictosidine-β-D-glucosidase; *GS*, geissoschizine synthase; *GO*, geissoschizine oxidase; *SAT*, stemmadenine-O-acetyltransferase; *PAS*, precondylocarpine acetate synthase; *DPAS*, dihydroprecondylocarpine synthase; *CS*, catharanthine synthase; *TS*, tabersonine synthase. The late pathway: *T16H1*, tabersonine 16-hydroxylase 1; *T16H*, tabersonine 16-hydroxylase 2; *16OMT*, tabersonine 16-O-methyltransferase; *T3O* - tabersonine 3-oxygenase; *T3R* - tabersonine 3-reductase; 16*NMT*, 16-methoxy-2,3-dihydro-3-hydroxytabersonine N-methyltransferase; *DAT*, deacetylvindoline-4-O-acetyltransferase; *D4H*, desacetoxyvindoline-4-hydroxylase; *PRX1*, class III peroxidase 1. IDIO+ gene IDs in Table S7.

**Table 1.** Top 25 upregulated genes in idioblast protoplasts, compared with protoplasts of common mesophyll cells, ranked by log2 fold change. Manually annotated genes from the MIA pathway are in bold, as well as gene families implicated in the MIA pathway. Transcripts per million (TPM) are the average of three idioblast protoplast samples.

| GeneID | Log2FC | Annotation | TPM |
| --- | --- | --- | --- |
| CATHA_25538 | 12.92 | Glycosyltransferase | 109.9 |
| CATHA_32840 | 11.73 | Aromatic amino acid decarboxylase 2 | 53.4 |
| CATHA_24235 | 11.28 | Small GTPase superfamily | 109.1 |
| CATHA_18760 | 10.92 | Absciscic stress-ripening protein 2 | 45.3 |
| CATHA_15400 | 10.90 | **2-Oxoglutarate (2OG) and Fe(II)-dependent oxygenase superfamily protein** | 43.9 |
| CATHA_18104 | 10.77 | Beta-amylase | 841.4 |
| CATHA_5278 | 10.23 | Sugar transporter | 18.4 |
| CATHA_18617 | 10.23 | **Deacetylvindoline-4-O-acetyltransferase (DAT)** | 65.1 |
| CATHA_12642 | 10.09 | **Tetrahydroalstonine synthase (THAS)** | 20.4 |
| CATHA_33484 | 10.01 | DUF679 domain membrane protein 2 | 695.5 |
| CATHA_10489 | 10.00 | **Serpentine synthase** | 75.4 |
| CATHA_38357 | 10.00 | Hypothetical protein | 293.2 |
| CATHA_33482 | 9.95 | DUF679 domain membrane protein 2 | 3725.2 |
| CATHA_30029 | 9.73 | Nodulin MtN21 /EamA-like transporter family protein (Fragment) | 55.1 |
| CATHA_26821 | 9.73 | **Nitrate transporter (NPF2)** | 989.6 |
| CATHA_29024 | 9.71 | Premnaspirodiene oxygenase-like (71D) | 12.7 |
| CATHA_20042 | 9.68 | Transcription repressor MYB4 | 612.8 |
| CATHA_33483 | 9.64 | DUF679 domain membrane protein 2 | 187.1 |
| CATHA_32017 | 9.60 | Major latex protein domain containing protein | 1547.4 |
| CATHA_5494 | 9.46 | Major latex | 580.6 |
| CATHA_22670 | 9.46 | Phytocyanin domain-containing protein | 214.6 |
| CATHA_21679 | 9.44 | **MATE efflux family protein** | 258.9 |
| CATHA_24905 | 9.42 | **WRKY transcription factor** | 145.3 |
| CATHA_27906 | 9.38 | Purine permease (PUP) | 334.7 |
| CATHA_27908 | 9.29 | Purine permease (PUP) | 710.1 |

Expression analysis of enzyme genes from MIA pathway branches other than the one leading to the anticancer VLB is shown in Supplementary Fig. S8, together with the analysis of transporters and transcription factors previously implicated in the MIA pathway. *THAS1/2*, vitrosamine synthase (*VAS*) and vincadifformine 19-hydroxylase (*V19H*) are significantly more expressed in idioblasts, as well as the ATP-binding cassette G (ABCG) transporter *CrTPT2,* thought to be responsible for the efflux of catharanthine to the leaf surface; however, the expression level of *CrTPT2* is very low. No other transporter or transcription factor implicated in the MIA pathway was upregulated in idioblasts.

### Differential gene expression reveals candidate genes to missing events of the MIA pathway

Alkaloid analysis (Fig. 2B) confirmed idioblasts as a central accumulating spot of *C. roseus* leaf alkaloids and the putative site of VLB and VCR biosynthesis. As such, DGE analysis was screened for candidate genes for VLB and VCR biosynthesis, for transport events necessary for MIA accumulation in idioblast cells and their vacuoles, and for transcription factors potentially regulating VLB biosynthesis and MIA metabolic flux.

#### Candidate enzymes

Idioblast upregulated genes belonging to gene families involved in the biosynthesis of specialised metabolites are shown in Fig. 6. Overall, 22 cytochromes P450 (P450s), 7 alcohol dehydrogenases (ADH), 3 short-chain dehydrogenases (SDR) and 7 class III peroxidases were identified, together with other enzyme genes such as berberine bridge enzymes, methyltransferases, amongst others. Several of those genes cluster in module 6 or module 9, further supporting their candidate nature to MIA biosynthetic reactions. Two alcohol dehydrogenases (CATHA_14789 and CATHA_31565) are similar to *Rauvolfia spp.* vomilenine reductase 2 (81% identity, 98% query coverage, at aminoacid level), which is involved in ajmaline biosynthesis (Geissler *et al.*, 2016). They also present similarity with *C. roseus* DPAS (67% identity 92%, query coverage, at aminoacid level). Both these genes cluster in module 6 (Fig. 4A), which is highly correlated with alkaloids levels, making them excellent candidates to missing MIA biosynthetic steps.

**Fig. 6.**
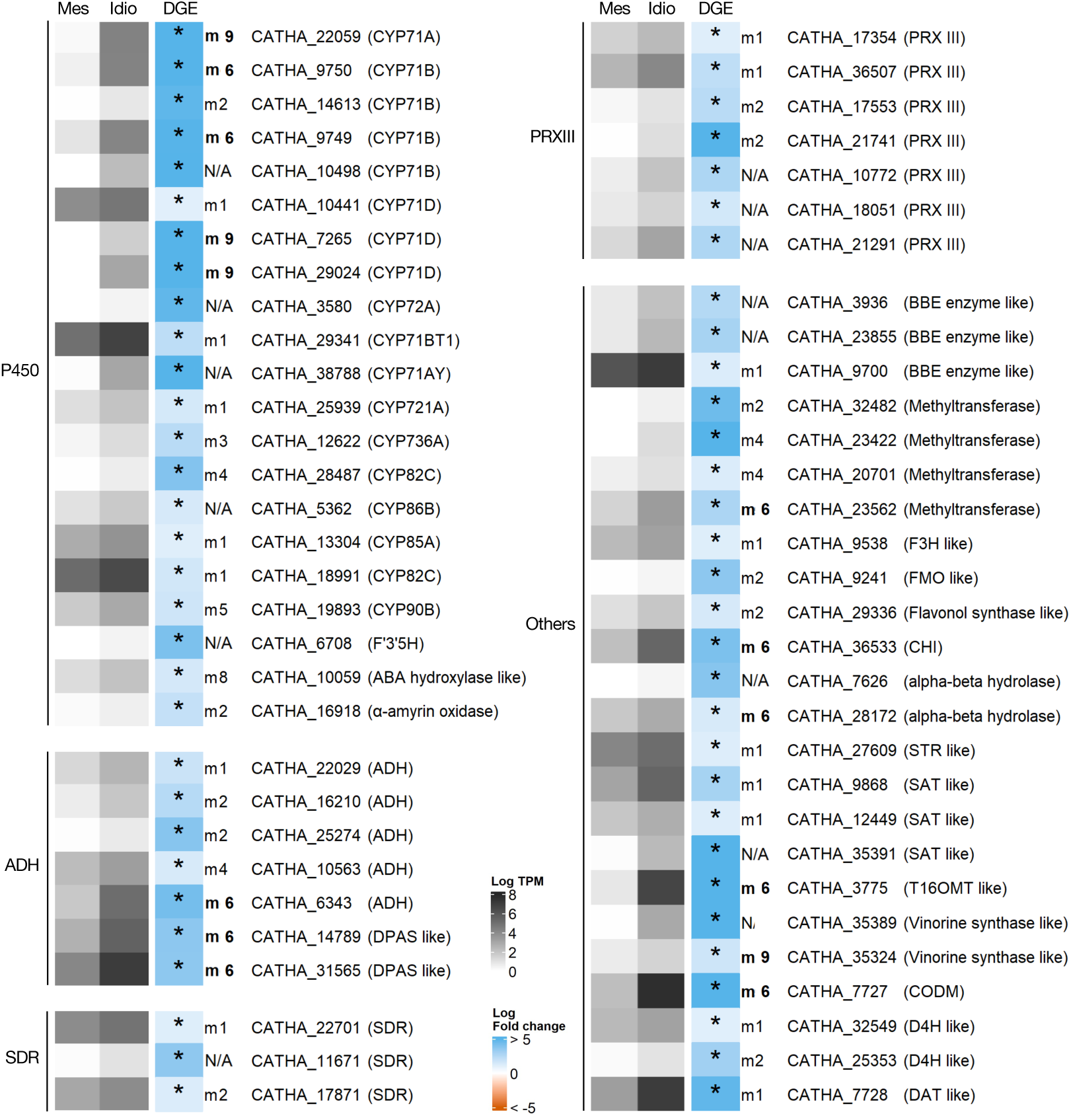
Expression analysis of candidate genes to MIA biosynthetic steps. P450 - cytochromes P450, ADH - alcohol dehydrogenases; SDR - short-chain dehydrogenases; PRX - class III peroxidases. Heatmap grey colours represent the mean log_2_ TPM of three independent biological samples. Log_2_ fold change for each gene is represented next to the expression level, with blue and red orange representing respectively overexpression and underexpression in idioblast protoplasts compared with common mesophyll protoplasts. Significant differential expression is marked with *. F3’5’H, flavonoid 3’5’ hydroxylase; ADH, alcohol dehydrogenase; DPAS, dihydroprecondylocarpine synthase; SDR, short-chain dehydrogenases; PRX III, class III peroxidase; BBE, berberine-bridge enzyme; F3H, flavanone 3-hydroxylase; FMO, flavin-containing monooxygenase; CHI, chalcone-flavanone isomerase; STR, strictosidine synthase; SAT, stemmadenine-O-acetyltransferase; T16OMT, tabersonine 16-O-methyltransferase; CODM, Codeine O-demethylase like; D4H, desacetoxyvindoline-4-hydroxylase; DAT, deacetylvindoline-4-O-acetyltransferase.

In order to further rank the P450s from the *C. roseus* idioblastome, a phylogenetic analysis was performed (Fig. 7A and Supplementary Fig. S9). The P450 family is the largest gene family in plants, with substrate specificity being somewhat phylogenetically conserved (Zheng *et al.*, 2019, Hamberger and Bak, 2013). Indeed, almost all P450s involved in MIA biosynthesis, as well as the P450 candidate genes from modules involved in alkaloid biosynthesis (modules 6 and 9) belong to clan 71. As such, phylogenetic inferences were conducted using only members of this subdivision, to which were added genes from other plant origins with known roles in specialized metabolism (Supplementary Table S10). To generate a robust phylogeny, analysis incorporated the complement of P450 clan 71 proteins from the eudicot and monocot model species *Arabidopsis thaliana* (AT) and *Oryza sativa* (Os), and of the angiosperm basal lineage species *Amborella trichopoda* (ATR). Fig. 7A shows that many of the idioblast P450 candidates align very close to P450 enzymes known to be involved in MIA biosynthesis in *C. roseus,* or in terpene metabolism in other species, and are therefore strong candidates for uncharacterized MIA reactions.

**Fig. 7.**
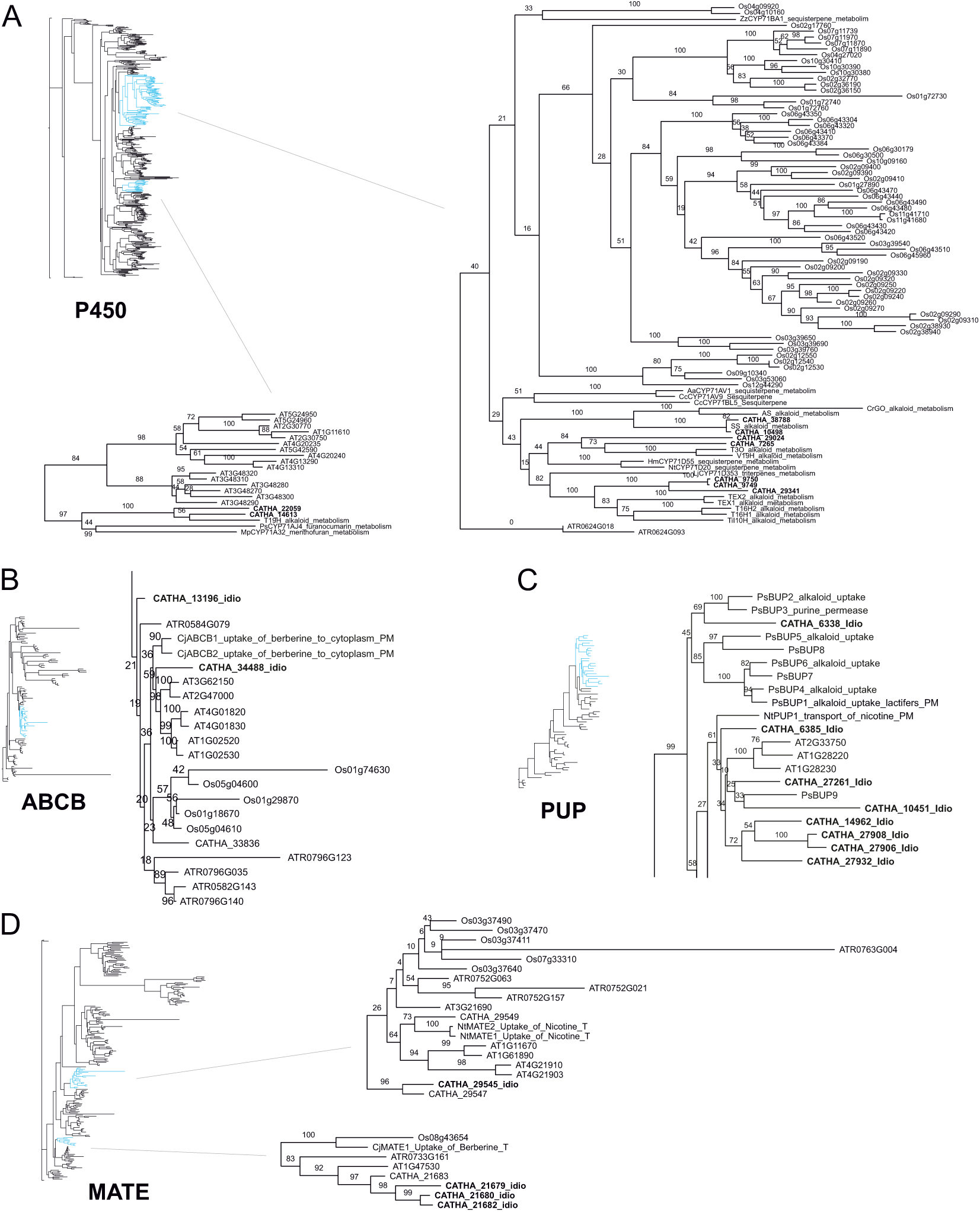
Relevant sections from the phylogenetic trees of *C. roseus* selected candidate proteins to MIA biosynthetic steps and MIA transmembrane transport, with the whole family proteins from several model species. (A) Cytochromes P450 from clan 71. (B) ABCB transporters. (C) PUP transporters. (D) MATE transporters. MATE proteins previously implicated in alkaloid transport appear in two independent clades. For each gene family tree, the phylogenetic analysis included the selected *C. roseus* proteins upregulated in idioblasts, the respective family proteins from *Arabidopsis thaliana* (AT), *Oryza sativa* (Os) and *Amborella trichopoda* (ATR) genomes, and the family proteins with known roles in specialised metabolism from other plant origins (Supplementary Tables S9-S12). The full trees can be found in Supplementary Figures S9-S12. Species initials: Aa, *Artemisia annua*; Cc, *Cynara cardunculus*; Cj, *Coptis japonica*; Hm, *Hyoscyamus muticus;* Lj, *Lotus japonicus*; Mp, *Mentha piperita*; Nt, *Nicotiana tabacum*; Ps (in C), *Papaver somniferum;* Ps (in A), *Pastinaca sativa;* Sm, *Salvia miltiorrhiza;* Ti, *Tabernanthe iboga*; Zz, *Zingiber zerumbet*. Enzymes (A): AS, alstonine synthase; I10H, ibogamine 10-hydroxylase; T3O, tabersonine 3-oxygenase; T16H, tabersonine 16-hydroxylase; TEX, tabersonine epoxidase; V19H, vincadifformine-19-hydroxylase.

#### Candidate transcription factors

Transcription factors belonging to gene families including members already implicated in the regulation of MIA metabolism in *C. roseus* were searched among upregulated genes in idioblasts. In total, 4 AP2-domain (apetala 2), 6 bHLH (basic helix-loop-helix), 9 WRKY genes and 19 MYB-domain (myeloblastosis) containing genes were retrieved (Fig. 8A). The full list of upregulated transcription factors in idioblasts is shown in Supplementary Table S11 and further includes several homeobox genes and plant specific families of transcription factors, such as NACs, GARPs and LOBs.

**Fig. 8.**
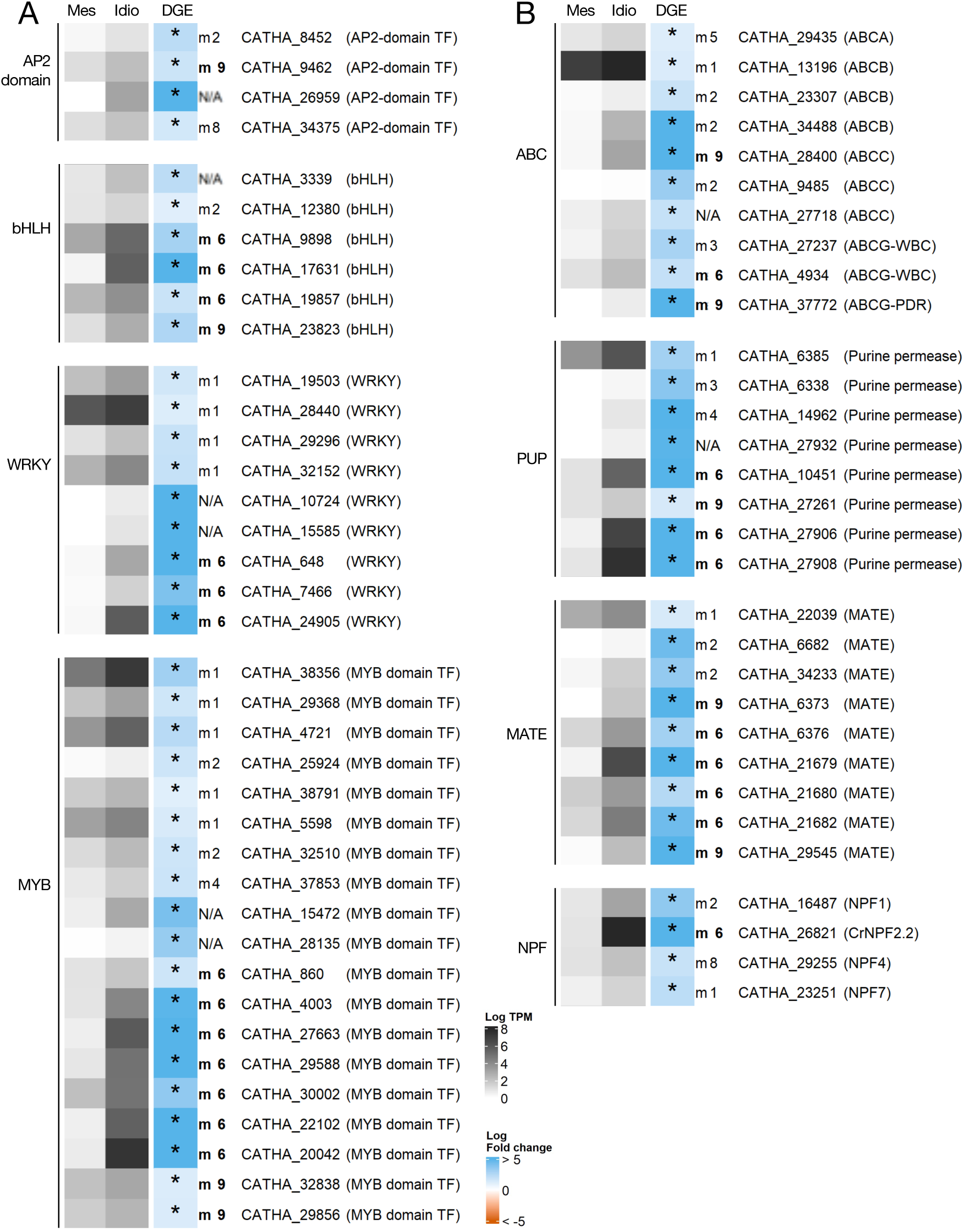
(A) Expression analysis of candidate genes to transcriptional regulation of the MIA pathway. AP2-domain TF, Apetala 2 domain transcription factor; bHLH, basic helix-loop-helix; MYB domain TF, myeloblastosis domain transcription factor. (B) Expression analysis of candidate genes to MIA transmembrane transport events. ABC, ATP-binding cassette transporter; PUP, purine permease; MATE, multidrug and toxic compound extrusion transporter; NPF, nitrate/peptide transporter. Heatmap grey colours represent the mean log_2_ TPM of three independent biological samples. Log_2_ fold change for each gene is represented next to the expression level, with blue and red orange representing respectively overexpression and underexpression in idioblast protoplasts compared with mesophyll protoplasts. Significant differential expression is marked with a *.

#### Candidate transporters

At least two major MIA transport events must occur in idioblasts: the transport of MIA intermediates from the apoplast to the cytosol, and the uptake into the vacuole of all the alkaloids accumulated in high levels in these cells (Fig. 2B). The list of genes upregulated in idioblasts was queried for members of transporter families known to have a role in transmembrane transport of specialised metabolites (Carqueijeiro *et al.*, 2013; Gani *et al.*, 2020). This search yielded several MIA candidates belonging to the transporter gene families MATE (9), ABC (10), PUP (8) and nitrate and peptide transporter family (NPF) (4) (Fig. 8B). In order to further rank the candidate nature of those genes, a phylogenetic analysis was performed, aligning the idioblast candidate proteins of each transporter family with proteins of the respective families previously implicated in transport of specialised metabolites, in *C. roseus* or other plant species (Fig. 7B-D, Supplementary Figs 10-12 and Suplementary Tables S12-S14). As for P450s, analyses incorporated the complement of family proteins from *A. thaliana* (AT), *O. sativa* (Os), and *A. trichopoda* (ATR). Fig. 7B-D shows that there are several upregulated idioblast proteins that are highly similar to known MATE, PUP, and ABCB transporters that were previously shown to be involved in alkaloid transmembrane transport, and are therefore strong candidates to the transmembrane transport of MIAs across the idioblast plasmalemma or tonoplast.

## Discussion

### The novel transcriptome IDIO+ represents an important upgrade for *C. roseus*

The importance of the anticancer alkaloids produced by *C. roseus* has driven the generation of multiple transcriptomic resources, which were pivotal for the full disclosure of the biosynthetic pathway of vindoline, catharanthine and serpentine (Gongora-Castillo *et al.*, 2012; Van Moerkercke *et al.*, 2013; Dugé de Bernonville *et al.*, 2015b; Franke et al. 2019). However, even if two improved genome versions have just been published (Cuello *et al.*, 2022; Sun *et al.* 2023), the transcriptome complexity still presents challenges namely concerning gene models and alternative splicing events.

Here, a novel transcriptome for *C. roseus*, IDIO+, was assembled using a genome-guided approach refined by Mikado (Supplementary Fig. S3). A major differentiation factor is that the RNA-seq data included transcript information from isolated idioblasts, which are low abundant cells in leaves, guaranteeing representation of otherwise rare transcripts. Thus, the likelihood of including undisclosed transcripts involved in the MIA pathway was substantially increased, given the high alkaloid content of idioblasts. The dataset also included data from plants grown outdoors, seldom used in research, and incorporated Nanopore long-read sequencing data, significantly enhancing the quality of the final gene-models.

IDIO+ is comparable, in terms of BUSCO scores, with the previously published consensus *de-novo C. roseus* transcriptome assembly CDF97 (Supplementary Table S7) (Dugé de Bernonville *et al.*, 2015b). However, the high number of gene-clusters in CDF97, as well as the high number of duplicated BUSCO orthologs indicates redundancy. Moreover, the building of CDF97 relied on CD-HIT-EST to cluster isoforms based on sequence similarity, which yields non-optimal results (Fu *et al.*, 2012; Davidson and Oshlack, 2014; Chen *et al.*, 2019). Indeed, *C. roseus* transcriptomic projects have underestimated the true number of paralogs for MIA pathway enzymes, due to high threshold being picked for isoform clustering, causing similar genes to be lost (Dugé de Bernonville *et al.*, 2015b). The use of genome-guided methods should circumvent these issues, as no clustering step is required.

IDIO+ improves the transcriptomic knowledge of *C. roseus* in several ways. The new RNA seq data allowed to uncover alternative splicing events and transcriptomic isoforms, adding new layers of information to the genome. Furthermore, 1,128 novel gene loci were identified (many corresponding to idioblast expressed genes) and there was an increase in the number of core orthologs found by BUSCO, in comparison with the genome v2 annotation (Supplementary Table S7). Finally, through the use of Mikado (Venturini *et al.*, 2018) and long-read data, it was possible to refine several erroneous reference gene-models, as exemplified in Supplementary Fig. S4. Overall, IDIO+ represents a novel asset to the species.

### The idioblastome reveals a unique cell type committed with defense

The term “idioblast” is used to describe individual cells that differ in shape, size, wall structure and/or content from a surrounding homogeneous tissue, and they may appear in any of the tissues of a plant (Foster, 1956). In *C. roseus,* the term was initially applied to isolated parenchyma cells identified by their specific reaction with alkaloid reagents (Yoder and Mahlberg, 1976). Later, it was shown that *C. roseus* leaf idioblasts also possess a distinctive autofluorescence, less chloroplasts, and larger size than surrounding mesophyll cells (Mersey and Cutler, 1986; Carqueijeiro *et al.*, 2016). In 1999, St. Pierre *et al.*, reported that the two last steps of vindoline biosynthesis (catalysed by DAT and D4H -Fig. 1) were specifically expressed in idioblasts and laticifers of *C. roseus* leaves. High differential expression of *DAT* and *D4H* was also detected in putative *C. roseus* leaf idioblast cells using single-cell cell RNA sequencing (Sun *et al.*, 2023).

The differential transcriptome of *C. roseus* leaf idioblasts generated in this work shows the expression of a clearly distinct genetic program from their surrounding mesophyll cells. Massive downregulation of genes involved in photosynthesis/light harvesting and chlorophyll binding (79% and 72% of genes associated with those GO terms, respectively; Fig. 3) and general downregulation of numerous gene categories related with photosynthesis and chloroplast processes / components indicate that idioblasts are not centrally engaged in light harvesting and carbon fixation, unlike common mesophyll cells. Actually, the strong upregulation of beta-amylase, of a sugar transporter and of beta-fructofuranosidase (Tables 1, 2) suggest that idioblasts are carbon sinks rather than carbon sources. Significantly, higher activity of beta-fructofuranosidase has been correlated with the utilisation and storage of sugars in sink organs, and with higher expression of specialised metabolism (Sturm, 1999; Nishanth *et al.*, 2018).

Correlated with the distinct behaviour concerning carbon metabolism, the GO enrichment analysis indicates that idioblasts show a radically different transport protein landscape (Fig. 3), certainly also related with their specialisation in alkaloid metabolism and accumulation (Fig. 2B). Accordingly, the top-ranking upregulated genes in idioblasts include, apart from the above-mentioned sugar transporter, several transporters from gene families involved in alkaloid transport discussed below as MIA candidates (MATE, PUP and NPF), and a bidirectional amino acid transporter (MtN21/EamA-like transporter) that may be important to feed idioblasts with aminoacid precursors. MtN21/EamA-like transporters have namely been implicated in glutamine transport (Denancé *et al.*, 2014) and its upregulation in idioblasts (∼10 log2 fold change, Table 1) may be related with the fact that glutamine synthetase is also upregulated (∼5 log2 fold change, Table 1), and among the top 25 genes with higher expression levels in idioblasts (Table 2). This indicates an intense nitrogen metabolism that may feed idioblasts with an abundance of aminoacids, including the MIA precursor tryptophan. Moreover, a high glutamine production was previously associated with ROS production and stress tolerance in Arabidopsis (Ji *et al.*, 2019).

**Table 2.** Top 25 upregulated genes in idioblast protoplasts, compared with protoplasts of common mesophyll cells, ranked by expression levels (TPM - transcripts per million). Manually annotated genes from the MIA pathway are in bold. TPMs are the average of three idioblast protoplast samples.

| GeneID | Log2FC | Annotation | TPM |
| --- | --- | --- | --- |
| CATHA_15668 | 5.86 | Major latex protein domain containing protein | 7199.5 |
| CATHA_22715 | 1.18 | 1-Aminocyclopropane-1-carboxylate oxidase | 5773.2 |
| CATHA_35751 | 1.83 | Beta-fructofuranosidase, insoluble isoenzyme CWINV1-like | 4103.5 |
| CATHA_10184 | 1.82 | Hypothetical protein | 4078.3 |
| CATHA_33482 | 9.95 | DUF679 domain membrane protein 2 | 3725.2 |
| CATHA_19823 | 3.39 | Dehydrin | 3393.2 |
| CATHA_16221 | 2.66 | Glutamic acid-rich protein-like isoform X1 | 3347.6 |
| CATHA_13610 | 1.78 | Dehydrin | 2585.9 |
| CATHA_7984 | 2.48 | Hexosyltransferase | 2382.5 |
| CATHA_9897 | 1.97 | Response to dessication 2 | 2107.2 |
| CATHA_15979 | 1.21 | J domain-containing protein | 2074.7 |
| CATHA_36137 | 1.21 | GDSL-like Lipase/Acylhydrolase superfamily protein | 1973 |
| CATHA_23202 | 2.88 | Histone deacetylase HDT1-like | 1964 |
| CATHA_9026 | 1.09 | Hypothetical protein | 1786.8 |
| CATHA_18758 | 2.62 | Similar to *Saccharomyces cerevisiae* YMR173W DDR48 DNA damage-responsive protein | 1770.5 |
| CATHA_32017 | 9.60 | Major latex protein domain containing protein | 1547.4 |
| CATHA_25357 | 4.53 | **Desacetoxyvindoline-4-hydroxylase (D4H)** | 1541.2 |
| CATHA_8113 | 2.45 | Polygalacturonase inhibitor | 1501.8 |
| CATHA_26149 | 7.39 | Patatin | 1386.8 |
| CATHA_12222 | 5.02 | Glutamine synthetase | 1354.1 |
| CATHA_29872 | 1.23 | Nematode resistance protein-like HSPRO2 | 1348.8 |
| CATHA_34966 | 1.17 | Malic enzyme | 1329.5 |
| CATHA_26934 | 1.84 | NAC domain-containing protein | 1300.1 |
| CATHA_18632 | 1.44 | Kunitz trypsin inhibitor 2-like | 1241.9 |
| CATHA_7086 | 5.69 | Isoflavone reductase homolog | 1233.8 |

Second to transmembrane transport, “response to biotic stimulus” and “response to oxidative stress” are the GO terms more represented in idioblast upregulated genes (Fig. 3). Accordingly, some of the genes with highest expression/upregulation in idioblasts are related with plant defence responses, including several major latex proteins (MLP), patatin, nematode resistance protein-like, isoflavone reductase, etc. (Tables 1, 2) (Lytle *et al.*, 2009; Cheng *et al.*, 2015; Bártová *et al.*, 2019). Notably, isoflavone reductases are found only in plants and they are considered to be very important to defence against several biotic stresses (Cheng *et al.*, 2015). Several of the top idioblast upregulated genes have also been implicated in abiotic stress, such as abcisic acid ripening 1 (*ASR1*), dehydrin and phytocianin (Golan *et al.*, 2014; Cao *et al.*, 2015). Therefore, the idioblast genetic program seems to be particularly fitted to respond to stress and most likely to sense stress. The signalling differential specialisation of idioblasts is suggested by the high number / “category %” of idioblast upregulated genes annotated with the GO terms “integral component of membrane” (557 genes) and “phosphatidylinositol phospholipase C activity” (57% genes of the category) (Fig. 3). The ∼11 log2 fold expression of a small GTPase, which is the third gene with higher log2 fold change (Table 1), also indicates that idioblasts are quite distinct from mesophyll cells in what concerns signalling, since small GTPases are molecular switches that typically function as network nodes integrating broad regulatory inputs into broad effector outputs (Reiner and Lundquist, 2018). A further indication of the higher capacity of idioblast cells to sense and respond to stress, is the fact that the GO terms “cell wall biogenesis”, “cell wall thickening” and “cell wall” are enriched in this cell type, suggesting that these cells respond more efficiently to the protoplasting aggression than common mesophyll cells. Consequently, idioblast cells are likely more responsive to cell wall damage resulting from herbivore attack.

All differential analyses indicate a strong specialisation of idioblasts in alkaloid metabolism. *D4H* and *DAT*, involved in the late biosynthesis of anticancer alkaloids (Fig. 1), and previously shown to be specifically expressed in idioblasts and laticifers by *in situ* hybridization and immunocytochemistry (St. Pierre *et al.*, 1999), appear among the top upregulated genes in idioblasts (Tables 1, 2). This confirms the previous results of St. Pierre *et al.* (1999), as well as the identity of the idioblast protoplast populations used in this study. The GO terms “alkaloid metabolic process” (23 genes, 21%) and “alkaloid biosynthetic process” (7 genes, 44%) appear overrepresented in idioblasts, while “alkaloid metabolic process” (23 genes, 21%) is also enriched in the downregulated genes. This indicates that idioblasts are specialised in specific parts of the MIA pathway, something that becomes clear when the behaviour of the genes involved in MIA biosynthesis is analysed (Fig. 5, discussed below).

*C. roseus* idioblasts accumulate high levels of alkaloids (Fig. 2B), which are stored inside the idioblast vacuole, avoiding self-toxicity effects (Carqueijeiro *et al.*, 2013). The singularity of idioblast vacuoles is further indicated by the enrichment of GO terms “plant-type vacuole” in both up and down regulated genes in idioblasts. Moreover, DUF679 domain membrane protein genes, which appear among the genes with highest log2 fold upregulation and expression levels in idioblasts (Tables 1, 2), have been implicated in tonoplast and ER membrane remodelling, namely during vacuole biogenesis (Kasaras *et al.*, 2012).

Finally, the major presence of three MLPs in idioblasts should be highlighted, with *CATHA_15668* corresponding to the gene with highest expression in this cell type (Table 2). MLPs are plant-specific proteins that were first identified as two of the most abundant proteins in the alkaloid accumulating latex of opium poppy (*Papaver somniferum* L.) (Nessler *et al.*, 1985), and they have been implicated in drought and salt tolerance, as well as in the induction of resistance against pathogens (Fujita and Inui, 2021). Their major presence in the alkaloid accumulating idioblasts of *C. roseus* clearly relates with a similar presence in the alkaloid accumulating poppy latex, and suggest a common defence role, most likely against herbivore attacks and subsequent pathogen exposure. The defense action mechanism of MLPs and their potential use in crop protection should be investigated.

Overall, the idioblastome reveals a unique cell type, clearly committed with defense against abiotic and biotic stresses. The overall functional portrait highlighted by the idioblastome, together with the high levels of alkaloids (Fig. 2B), whose animal toxicity and bitterness guarantees potent herbivore-deterrent activity, indicate that *C. roseus* leaf idioblasts must be instrumental in responses to and resistance against environmental challenges. In the plant kingdom, there are several examples where such individual cells play an important role in defense mechanisms, with the Brassicacea myrosin cells being one of the most well studied examples (Rodriguez-Saona and Trumble, 2000; Shirakawa and Hara-Nishimura, 2018; Eco and Belonias, 2017).

### A new paradigm for alkaloid accumulation in *C. roseus* leaves

In this work, analysis of the alkaloid profile of idioblast protoplasts revealed the differential presence of extremely high levels of serpentine, catharanthine, vindoline and AVLB, which were also the main alkaloids detected in *C. roseus* leaves (Fig. 2B). The relative proportions of the different alkaloids in idioblast protoplasts were very similar to the ones observed in leaves and total protoplasts, suggesting that the alkaloid content of idioblasts is making a decisive contribution to the total alkaloid content of leaves. These results challenge the current view of alkaloid accumulation in *C. roseus* leaves, which predicts that catharanthine is accumulated at the epidermal cuticle, away from the idioblast-accumulated vindoline (original data: Roepke *et al.*, 2010; Yu and De Luca, 2013; important reviews: De Luca *et al.*, 2014; Courdavault *et al.*, 2014; Dugé de Bernonville *et al.*, 2015a; Kulagina *et al.*, 2022). Here, the observed matching of the alkaloid profile of idioblasts with that of leaves is incompatible with any significant accumulation of catharanthine in the epidermis or cuticle of the plants used. Furthermore, the presence of high levels of AVLB indicates the occurrence of intense dimerization, with co-localization of catharanthine and vindoline. It should be emphasised that the idioblast protoplasts obtained with the methodology used here were previously shown to retain their *in planta* distinctive identity (Carqueijeiro *et al.*, 2016).

In fact, the results of this study are in line with abundant published evidence that has been overlooked. The presence of high levels of AVLB in *C. roseus* leaves has been reported at least by Balsevich and Bishop (1989), Naaranlahti *et al.* (1991), Tang *et al.* (2009), Pan *et al.* (2016), and in our lab, by Sottomayor *et al.* (1998), Costa *et al.* (2008) and Carqueijeiro *et al.* (2013), spanning at least three different cultivars, across six distinct geographical locations. Carqueijeiro *et al.*, 2013 showed that mesophyll protoplasts accumulate high levels of vindoline, catharanthine and AVLB, indicating cellular accumulation of catharanthine and co-localization with vindoline to yield AVLB. Furthermore, it was shown that *C. roseus* mesophyll vacuoles perform specific uptake of all three alkaloids through a H^+^-antiport system, reinforcing the evidence supporting cellular accumulation of catharanthine. Mersey and Cutler (1986) also reported an enrichment of catharanthine and vindoline in a density fraction enriched in leaf idioblast protoplasts, while Yamamoto *et al.*, (2019) clearly showed by imaging and single-cell MS analysis that catharanthine accumulates in leaf idioblasts rather than in any other leaf location. Similarly, the treatment of *C. roseus* leaves with alkaloid staining reagents has consistently showed reaction with idioblast and laticifers, but never with the epidermis or the leaf surface (Yoder and Mahlberg, 1976; Carqueijeiro *et al.*, 2016; Usaki *et al.*, 2022).

The specific accumulation of catharanthine in the leaf surface was reported only by Roepke *et al.* (2010) and Yu and De Luca (2013) based on a leaf chloroform dipping technique that was claimed to show that catharanthine occurs exclusively on the surface of *C. roseus* leaves. However, Murata and De Luca (2005) from the same lab, using an abrasion technique to separate the epidermis from the leaf body, had previously reported that vindoline and catharanthine were both accumulated in the central part of the leaf body and not in the epidermis. Finally, Abouzeid *et al.* (2019) repeated the leaf chloroform dipping technique and demonstrated that chloroform destroys the cell integrity of *C. roseus* leaves, concluding that alkaloids are located exclusively in the *C. roseus* leaf symplast.

Therefore, there is a consistent body of evidence indicating that the accumulation pattern of alkaloids in *C. roseus* leaves involves mainly idioblasts and laticifers, with co-localization of the monomeric precursors catharanthine and vindoline in those cell types. As such, the low levels of the dimeric anticancer alkaloids VLB and VCR should not be credited to the spatial seclusion of their monomeric precursors catharanthine and vindoline, as often stated in recent literature (De Luca *et al.*, 2014; Courdavault *et al.*, 2014; Qu *et al.*, 2019; Kulagina *et al.*, 2022). Most studies show that *C. roseus* leaves accumulate high levels of catharanthine and vindoline, with high levels of AVLB also being frequently reported (see above). In these cases, the true bottleneck in what concerns the accumulation of the anticancer alkaloids is the conversion of AVLB into VLB, a reaction that has escaped characterization so far. There are other reports where AVLB levels are low, undetected or not mentioned, indicating that the dimerization reaction may also be an important bottleneck for specific *C. roseus* cultivars / growing conditions (e.g. Mersey and Cutler, 1986; Murata and De Luca, 2005; Dugé de Bernonville *et al.*, 2017; Roepke *et al.*, 2010; Yu and de Luca, 2013; Yamamoto *et al.*, 2019). In the plants used here, the dimerization activity is robustly co-regulated with the biosynthesis of the monomeric precursors (Fig. 4A), while VLB biosynthesis shows a strikingly different pattern of correlation with gene co-expression modules, indicating a completely distinct regulation from the accumulation of its direct precursors that may explain its elusive nature.

This study, together with the previous reports of Yoder and Mahlberg (1976), Mersey and Cutler (1986), Carqueijeiro *et al.* (2016), Yamamoto *et al.* (2019) and Usaki *et al.* (2022), establish idioblasts as a central accumulation target of alkaloids in *C. roseus* leaves, and rationally the home of the key biosynthetic steps leading to the dimeric anticancer alkaloids, of transporters determining alkaloid metabolic flux, and of instrumental transcriptional regulators of the MIA pathway (Fig. 9). Based on their similar fluorescent properties and reactivity with alkaloid reagents (Carqueijeiro *et al.*, 2016), laticifers are most likely a very similar alkaloid hot spot.

**Fig. 9.**
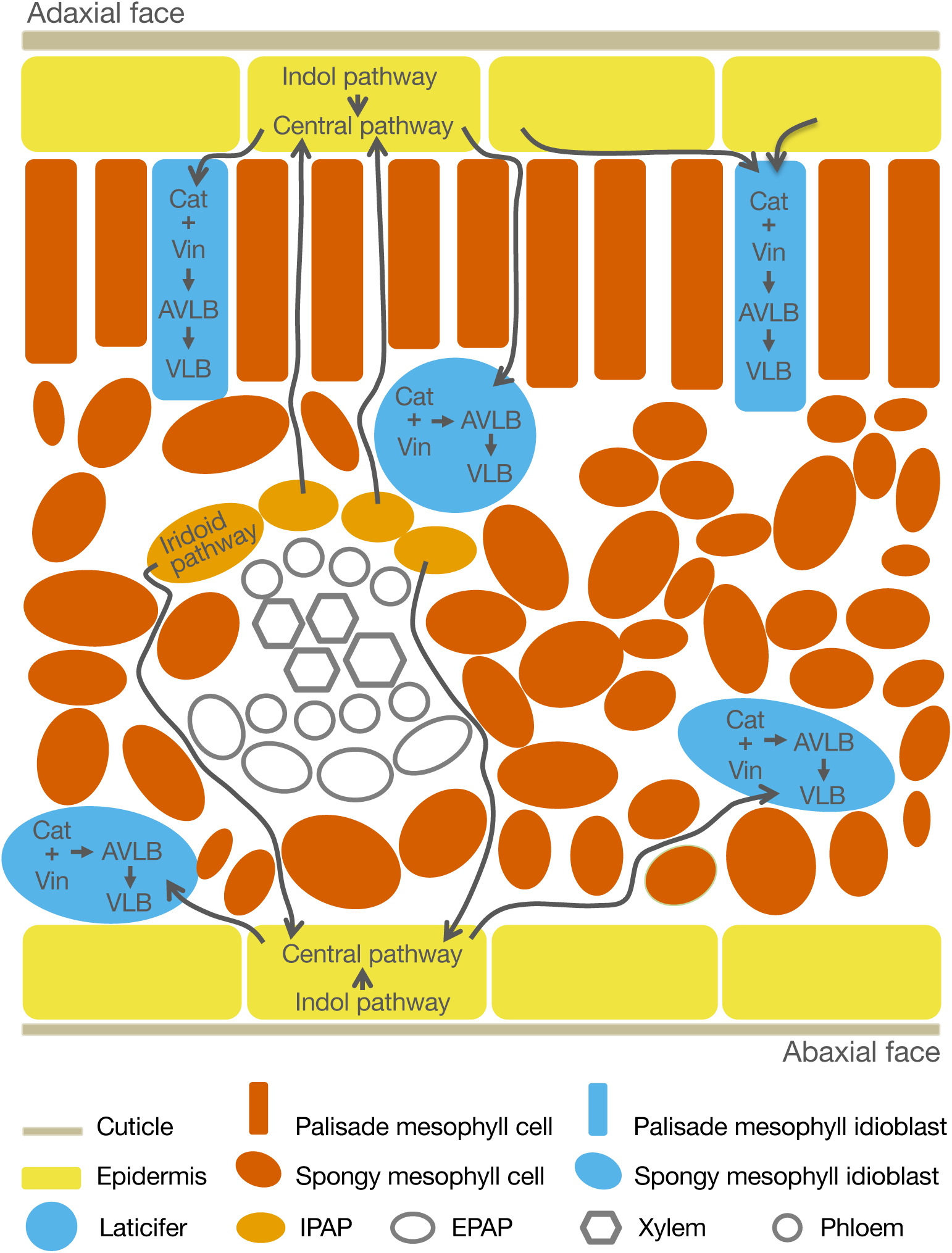
Cellular localization of the different parts of the MIA pathway in *C. roseus* leaves. Iridoid precursors are produced in IPAP cells and transported to epidermal cells where the central pathway takes place, also incorporating indole precursors. Catharanthine and a vindoline precursor are transported to mesophyll idioblasts and laticifers, where those monomeric alkaloids accumulate in high levels. The dimerization into AVLB and the subsequent conversions to VLB and VCR, the true bottlenecks of the pathway, also take place in idioblasts and laticifers. IPAP – internal phloem associated parenchyma. EPAP – external phloem associated parenchyma. For pathway details see Fig. 1.

This confinement of specialized metabolism to particular cell types is a recurrent feature in plants, with examples including morphine in poppy laticifers, glucosinolate metabolism in Arabidopsis myrosin cells and S-cells, and terpenoids in glandular trichomes of ∼30% of vascular plants (Glas *et al.*, 2012; Beaudoin and Facchini, 2014; Shirakawa and Hara-Nishimura, 2018). Such compartmentation, together with vacuole accumulation of most specialized metabolites, including MIAs, has been considered a strategy to manage the burdens of overproduction of specialized metabolites, including competition for resources, metabolic imbalance, and potential self-intoxication (Carqueijeiro *et al.*, 2013; Chezem and Clay, 2016).

### The idioblastome reveals new insights into the MIA pathway and establishes a roadmap towards its full elucidation

#### MIA biosynthesis

The differential expression of MIA biosynthetic genes in idioblasts shows that the indole, iridoid and central MIA pathways are downregulated in this cell type, while most of the late pathway, encompassing the conversion of tabersonine into vindoline is upregulated (Figs 1, 5). It is also evident that three of the branching pathways competing with the central pathway for the common strictosidine aglycone substrates are upregulated in idioblasts (Fig. 1, Supplementary Fig. S8). How the plant regulates the channelling of strictosidine aglycone into the divergent structural classes stemming from this central precursor is precisely one of the main riddles of the MIA pathway. Metabolic channeling resulting from interaction between strictosidine-β-D-glucosidase (SGD) and subsequent enzymes is probably one of the mechanisms involved, since it has been shown by bimolecular fluorescence complementation (BiFC) that heteroyohimbine synthase (HYS) and tetrahydroalstonine synthases (THAS1 and 2) interact *in vivo* with the enzyme producing strictosidine aglycone, SGD (Stravinides *et al.*, 2015, 2016). Our results suggest that differential subcellular localization may also be important to determine the fate of strictosidine aglycone, since geissochizine synthase 1 (GS1) transcripts, feeding the central pathway towards catharanthine and tabersonine, have been shown to be enriched in epidermal extracts (Tatsis *et al.*, 2017; Qu *et al.*, 2018a), while the competing THAS1/2 and vitrosamine synthase (VAS) are here shown to have differential higher expression in idioblasts.

Interestingly, heteroyohimbine synthase (HYS), the putative entry enzyme towards serpentine, an alkaloid present in high levels in idioblasts, is not differentially expressed in these cells, presenting very low absolute expression levels. This is the contrary of serpentine synthase (SS), which is expressed in differential and absolute high levels in idioblasts (Table 1 and Supplementary Fig. S9), in accordance with the observed high serpentine levels in those cells (Fig. 2B). It has been shown that silencing *THAS1* in *C. roseus* leaves induces a significant decrease in ajmalicine levels (Stravinides *et al.*, 2016). This fact, together with the idioblast high levels of THAS1, SS and serpentine observed here, suggest that THAS1 may be the enzyme acting together with SS to produce serpentine in idioblasts. Alternatively, ajmalicine could be produced elsewhere in the leaves, and then transported into idioblasts. However, heteroyohimbine synthase (HYS) was not found in epidermis-enriched extracts (Qu *et al.*, 2018a), and has low expression levels in mesophyll cells (Supplementary Fig. S8).

Strictosidine aglycone isomers are very unstable and toxic and it has always been assumed that they are promptly converted *in loco* by subsequent enzymes (Guirimand *et al.*, 2010; Stravinides *et al.*, 2016). The high expression levels of THAS1/2, vitrosamine synthase (VAS) and SS in idioblasts, together with the high levels of the alkaloid product serpentine, suggest that the entry enzymes of the MIA central pathway, strictosidine synthase (STR) and strictosidine-β-D-glucosidase (SGD), although downregulated in idioblasts, are also operating in this cell type.

An intriguing observation is the upregulation of *DPAS* in idioblasts, opposite to the immediately previous and subsequent enzymes in the pathway (Figs 1, 5). This enzyme has been proposed to generate the unstable intermediate dihydroprecondylocarpine acetate (Caputi *et al.*, 2018), which also acts as a hub substrate originating at least three different products (Fig. 1). *DPAS* is also the only enzymatic step of the MIA pathway found in co-expression module 6 (Supplementary Table S7), which is the only module showing a significant positive correlation with the levels of the main alkaloids accumulated in *C. roseus* leaves / idioblasts (Fig. 4A). DPAS may therefore be a key step influencing decisively the metabolic flux into the production of catharanthine and vindoline, and deserves further attention. Two alcohol dehydrogenases (*CATHA_14789* and *CATHA_31565*) presenting similarity with *DPAS* are also highly expressed in idioblasts (Fig. 6). These genes cluster in module 6 (Fig. 4A), they are similar to *Rauvolfia spp.* vomilenine reductase 2, which is involved in ajmaline biosynthesis (Geissler *et al.*, 2016), and their relevance for the MIA pathway should be investigated.

Another unexpected result is the upregulation in idioblasts of several genes involved in the conversion of tabersonine into vindoline that have been previously shown to be specifically or preferentially expressed in the epidermis (Guirimand *et al.*, 2011b; Qu *et al.*, 2015). Moreover, that upregulation is not continuous, since it is not observed for T3O and T3R, the 3rd and 4th of the 7 biosynthetic steps (Fig. 1, 5). This may indicate that the developed leaves used in this study may have a spatial organisation of the MIA pathway that is different from the young or very young leaves used in the previous studies (Guirimand *et al.*, 2011b; Qu *et al.*, 2015). It could also indicate that there may be undisclosed alternative T3O and T3R operating in idioblasts. Further studies will be needed to clarify these questions.

*PRX1*, a class III peroxidase putatively responsible for the coupling reaction of catharanthine and vindoline to yield the first dimeric MIA was downregulated in idioblasts. *PRX1* was also not co-expressed with *D4H* and *DAT*, in module 9, nor clustered in module 6, which has a high correlation with AVLB levels in the co-expression analysis (Fig. 4A). Alike to other class III plant peroxidases, evidence suggests that PRX1 is not a specific enzyme and has a wide range of roles, including scavenging of excess hydrogen peroxide generated during photosynthesis at the expense of oxidation of a range of different substrates (Ferreres *et al.*, 2011). Most photosynthetic and chloroplast processes are greatly downregulated in idioblasts (Fig. 3), what could explain the lower expression of *PRX1* in such cells, independently of its function as the dimerizing enzyme. Otherwise, it is not impossible that although PRX1 is capable of performing the coupling reaction *in vitro*, there is a more specific enzyme actually performing that reaction *in vivo,* in idioblasts. Moreover, the enzyme or factor responsible for the reduction to AVLB of the dimeric iminum ion generated by PRX1 has never been disclosed.

A striking outcome of module-trait correlation analysis is the radically different correlation behaviour of VLB in comparison with all their immediate precursor alkaloids (Fig. 4A). This indicates that the unknown enzyme responsible for the reaction leading to this valuable anticancer alkaloid might not be co-expressed with the other MIA pathway steps occurring in idioblasts. Moreover, no significant correlation was found between VLB and any of the gene co-expression modules, in line with its elusive biosynthesis.

The answer to the gaps in the knowledge about the last bottleneck biosynthetic steps leading to the anticancer VLB and VCR should be among the candidate enzymes of Fig. 6, featuring the two DPAS-like alcohol dehydrogenases mentioned above and the versatile cytochrome P450s. In fact, most of the idioblast P450 candidates are close to other proteins involved in specialized metabolism (Fig. 7A): CATHA_10498 and CATHA_38788 are close to alstonine synthase (AS) and *C. roseus* geissoschizine oxidase; CATHA_29024 and CATHA_7265 are close to vincadifformine 19-hydroxylase (V19H) and T3O; CATHA_9749, CATHA_9750 and CATHA_29341 are close to tabersonine epoxidase 1/2 (TEX1/2) and tabersonine 16-hydroxylase 1/2 (T16H1/2); and CATHA_14613 and CATHA_22059 are in the same subclade as tabersonine 19-hydroxylase (T19H). Additionally, several of these idioblast P450 genes cluster in the co-expression network either in module 6 or module 9 (Fig. 4A), reinforcing a potential role in the MIA pathway.

#### MIA transcriptional regulation

There seems to be a significant overlap between the transcriptional regulatory networks governing specialized metabolism and the development of associated cell types (Chezem and Clay, 2016). As concluded above, our data indicate that *C. roseus* mesophyll idioblasts are specialized in the late steps of the MIA pathway and centralize MIA accumulation in *C. roseus* leaves. Therefore, the transcription regulatory network determining idioblast differentiation and the maintenance of their differentiation status is most likely key for the regulation of the late MIA pathway and MIA metabolic flux. Among the up-regulated genes in idioblasts, 18 genes are annotated as involved in differentiation or polarity (Fig. 3, UP, BP), including three MYBs listed among candidate transcription factors: *CATHA_22102, CATHA_29588* and *CATHA_30002* (Fig. 8A). These MYBs are included in module 6, associated with alkaloid levels, and their biological process GO term annotation includes “cell differentiation” as the unique term. This suggests a possible central role in the regulation of the differentiation status of idioblasts and of their specific features concerning alkaloid metabolism. Several other candidate transcription factors clustered in module 6 or 9, including two other MYBs, three WRKYs and one bHLH (Fig. 8A), all from transcription factor families previously implicated in the regulation of the MIA pathway. They are therefore interesting candidates to the transcriptional regulation of the late MIA biosynthetic steps and metabolic flux.

#### MIA transmembrane transport

Correlation analysis of gene co-expression modules with alkaloid traits showed that only module 6 has a significant correlation with alkaloid levels (Fig. 4A). This module includes mostly genes upregulated in idioblasts, it does not include any MIA biosynthetic gene and it includes many transporter genes (Supplementary Table S9). This result indicates the relevance of idioblasts for alkaloid accumulation in *C. roseus* leaves and suggest that alkaloid levels are not determined by the expression levels of biosynthetic enzymes, but rather by the intensity of metabolic flux generated by alkaloid transmembrane transport. As such, the transporter candidates identified in the idioblast complement of genes (Fig. 8B) hold an exceptional potential in what concerns the understanding and improvement of alkaloid metabolic flux in *C. roseus* leaves. Significantly, module 6 includes several candidates with close phylogenetic relationships with transporters previously shown to be involved in alkaloid transport in other species (Fig. 7B-D, Fig. 8B).

ABC transporters have been implicated in the transmembrane transport of specialized metabolites, including members from the subfamilies ABCB, ABCC and ABCG (Gani *et al.*, 2020). In total, 10 ABC transporters were upregulated in idioblasts (Fig. 8B). Phylogenetic analysis of ABCBs revealed that CATHA_34488 clustered with CjABCB1 and CjABCB2, involved in alkaloid cellular uptake in *Coptis japonica* (Shitan *et al.*, 2003; Shitan *et al.*, 2013) (Fig. 7B). *CATHA_34488*, as all other ABCB candidates, did not cluster in module 6 or 9, but its relative proximity with *Coptis japonica ABC1/2* involved in berberine transport is suggestive of alkaloid transport.

From NPF transporter candidate genes (Fig. 8B), only one belongs to subfamily 2 (*CATHA_26821*), which is the only clade with known genes involved in the transport of specialised metabolites (Nour-Eldin *et al.*, 2012; Larsen *et al.*, 2017), including CrNPF2.9, which has been shown to export strictosidine from the vacuoles (Payne *et al.*, 2017). *CATHA_26821* is among the top 25 upregulated genes in idioblasts ranked by log2 fold change (Table 2), it has high expression levels in idioblasts and clusters in module 6. BLAST search revealed that CATHA_26821 corresponds to a previously studied *C. roseus* NPF2.2 transporter which was evaluated with negative results for the transport of MIA monoterpenoid precursors, but was not evaluated in what concerns MIAs themselves (Larsen *et al.*, 2017). In the same study, three other CrNPF2 transporters were shown to be localized in the plasma membrane. Therefore, the potential role of NPF2.2 in the idioblast import of MIA intermediates should be investigated.

Members of the purine uptake permease (PUP) transporters family were implicated in alkaloid transport from the apoplast to the cytosol, in *Nicotiana tabacum* and *Papaver somniferum* (Jelesko, 2012; Dastmalchi *et al.*, 2019). All the eight candidate PUP transporters upregulated in idioblasts (Fig. 8B) align in one strongly supported clade (Fig. 7C) containing the previously characterised alkaloid transporters, thus revealing a common evolutionary origin that may have functional significance. In *P. somniferum,* BUP1 acts as a generalist transporter for benzylisoquinoline alkaloids, while BUP2-6 displays preference for different pathway intermediaries (Dastmalchi *et al.*, 2019). It is therefore possible that the *C. roseus* PUP candidate transporters unveiled in this work have distinct alkaloid substrates. Three of the *C. roseus* PUPs cluster in module 6 of the co-expression network (*CATHA_10451, CATHA_27906, CATHA_27908*), while one clusters in module 9 (*CATHA_27932*), further strengthening their potential as candidates for MIA transport.

MATE transporters have been implicated in vacuole uptake of alkaloids, with NtMATE1/2 and CjMATE1 acting as nicotine/H^+^ and berberine/H^+^ antiporters in *Nicotiana tabacum* and *Coptis japonica* respectively (Otani *et al.*, 2005; Shoji *et al.*, 2009). In *C. roseus*, it has been shown that catharanthine, vindoline and AVLB are transported from the cytosol to the vacuole by a H^+^-antiport system evocative of MATE transporters (Carqueijeiro *et al.*, 2013). There were nine *CrMATES* upregulated in idioblasts (Fig. 8B), and phylogenetic analysis (Fig. 7D) showed that CATHA_29545 aligns in the clade of NtMATE1/2 and that CATHA_21679, CATHA_21680, and CATHA_21682 align in the clade of CjMATE1. *CATHA_29545* clusters in module 9, which includes *DAT* and *D4H*, while *CATHA_21679*, *CATHA_21680*, and *CATHA_21682* cluster in module 6, significantly correlated with alkaloid levels of alkaloids (Fig. 4A). Furthermore, *CATHA_21679* is among the top upregulated genes in idioblasts (Table 2). These are therefore very strong candidates to the idioblast vacuolar uptake of the different MIAs, the transport event that is potentially the main determinant of MIA metabolic flux in *C. roseus* leaves. The genes of the three CrMATEs clustering together in the phylogenetic tree are located very closely in the genome, suggesting a recent origin by tandem gene duplication that could be part of the evolutionary path of *C. roseus* towards specialization / neofunctionalization in the MIA pathway.

Given the exclusive accumulation of high levels of alkaloids in *C. roseus* leaf idioblasts (together with laticifers) it seems obvious that this cell type acts as a sink of whatever alkaloids are produced elsewhere in the leaves. On the other hand, the idioblast expression profile shows that these cells are particularly active in what concerns the last steps of biosynthesis of serpentine and vindoline and are the specific home of the critical biosynthetic steps leading to the dimeric anticancer alkaloids from the monomeric precursors vindoline and catharanthine. Therefore, the idioblastome holds the key to the identification of the genes codifying the enzymes responsible for the bottleneck biosynthetic steps leading to the anticancer VLB and VCR, the transporters determining alkaloid metabolic flux, and the transcription factors regulating the expression levels of both MIA enzyme and transporter genes. The present work provides a comprehensive roadmap towards the deep understanding of the MIA pathway twists and the future implementation of successful strategies to increase the levels of the *C. roseus* anticancer alkaloids.

##### Supplementary Data

The following supplementary data are available at JXB online.

Dataset S1. GTF file with the IDIO+ full mRNA gene models (including all alternative splicing isoforms) and the respective localization coordinates in *C. roseus* genome v2. This was the file used for DGE.

Dataset S2. FASTA file including IDIO+ mRNA coding sequences. Dataset S3. TSV file including the functional annotation of IDIO+.

Dataset S4. CSV file including the TPM expression levels of IDIO+ genes in the samples used for DGE.

Fig. S1. Principal Component Analysis (PCA) of all the Illumina short-read transcriptomic datasets (Fig. 2C).

Fig. S2. Scale-free topological indices at various soft-thresholding powers for the co-expression network used in the module-trait correlation analysis of Fig. 4A.

Fig. S3. Summary of the approach used for genome-guided transcriptome assembly.

Fig. S4. Screenshot obtained with the genome browser Jbrowse (jbrowse.org) depicting an example of an erroneous gene-model of a MATE protein present in the reference genome annotation (CRO_T123311).

Fig. S5. Principal component analysis (PCA) of the transcriptomic datasets generated for the different tissue and cell samples involved in the isolation of idioblasts (Fig. 2C, upper part).

Fig. S6. Hierarchical clustering of the different tissue and cell samples involved in the isolation of idioblasts, based on the regularized log transformation of normalized read counts per gene.

Fig. S7. Levels of the monomeric alkaloid precursors catharanthine + vindoline (A) and of the dimeric alkaloids AVLB + VLB + VCR (B) detected in leaves from plants growing indoors (in) and outdoors (out).

Fig. S8. Expression analysis of genes that have been implicated in MIA branching pathways (Fig. 1), in MIA transmembrane transport and in transcriptional regulation of the MIA pathway.

Fig. S9. Phylogenetic analysis of cytochrome P450s from clan 71 upregulated in C. roseus idioblasts.

Fig. S10. Phylogenetic analysis of ABCB transmembrane transporters upregulated in C. roseus idioblasts.

Fig. S11. Phylogenetic analysis of PUP transmembrane transporters upregulated in C. roseus idioblasts.

Fig. S12. Phylogenetic analysis of MATE transmembrane transporters upregulated in C. roseus idioblasts.

Table S1. List of novel genes identified by IDIO+.

Table S2. Genes upregulated in idioblast protoplasts in comparison with protoplasts of common mesophyll cells.

Table S3. Genes downregulated in idioblast protoplasts in comparison with protoplasts of common mesophyll cells.

Table S4. Genes not significantly differentially expressed in idioblast protoplasts in comparison with protoplasts of common mesophyll cells.

Table S5. Best fitting evolutionary model used for the phylogenetic analyses performed for each gene family.

Table S6. Overview of the sequencing data used for transcriptome assembly.

Table S7. Comparison of the IDIO+ transcriptome assembly with the *C. roseus* genome v2 annotation and the consensus CDF97 transcriptome.

Table S8. Manual annotation of the MIA pathway genes in the IDIO+ transcriptome.

Table S9. Gene lists of co-expression modules of Fig. 4.

Table S10. Proteins involved in specialized metabolism used in the phylogenetic analysis of the P450 family.

Table S11. Full list of putative transcription factors upregulated in idioblasts and candidate to regulation of MIA metabolism.

Table S12. Proteins involved in specialized metabolism used in the phylogenetic analysis of the ABCB family.

Table S13. Proteins involved in specialized metabolism used in the phylogenetic analysis of the PUP family.

Table S14. Proteins involved in specialized metabolism used in the phylogenetic analysis of the MATE family.

## Abbreviations

AVLB: α-3’,4’-anhydrovinblastine
BUSCO: benchmarking set of universal single-copy orthologs
D4H: desacetoxyvindoline-4-hydroxylase
DAT: deacetylvindoline-4-O-acetyltransferase
DGE: differential gene expression
DPAS: dihydroprecondylocarpine synthase
FACS: fluorescence-activated cell sorting
GO: gene ontology
MATE: multidrug and toxic compound extrusion
MIA: monoterpenoid indole alkaloid
NPF: nitrate and peptide transporter family
P450: cytochrome P450
PCA: principal component analysis
PRX1: class III peroxidase 1
PUP: purine permease
RT: retention time
T3O: tabersonine 3-oxygenase
T3R: tabersonine 3-reductase
THAS: tetrahydroalstonine synthase
TPM: transcript count per million
SS: serpentine synthase
VCR: vincristine
VLB: vinblastine

Note: For the sake of easy identification and correspondence with Fig. 1, abbreviations of enzymes of the alkaloid pathway are always given, even if not used afterwards in the isolated form (since not repeated more than three times).

## Acknowledgements

We would like to thank Sílvia Maia from Laboratório de Análise Estrutural – LAE, Centro de Materiais da Universidade do Porto for technical assistance with HPLC-Electrospray Ionization-Tandem Mass Spectrometry.

## Author contributions

JGG: conceptualization; wet lab experiments; FACS; Illumina RNAseq; initial transcriptome assembly (TA), DGE and GO analyses; biological analysis of data; writing (final draft, review and editing). RR: conceptualization; Nanopore RNAseq; final TA, DGE and GO analyses; co-expression-trait and phylogenetic analyses; biological analysis of data; writing (initial draft, review and editing). IC: conceptualization; optimization of wet lab methodologies and FACS; writing (review and editing). ALG: optimization of wet lab methodologies and FACS. CB: FACS. JA: conceptualization; supervision of Illumina RNAseq, initial TA and DGE analyses; writing (review and editing).HA: supervision of phylogenetic analyses; biological analysis of data; writing (review and editing).NAF: conceptualization; supervision of Nanopore RNAseq, final TA, DGE and GO analyses, and of co-expression-trait analysis; writing (supervision of initial draft, review and editing).MS: conceptualization, FACS, supervision and coordination at all levels, biological analysis of data, writing (supervision of initial draft, final draft, review and editing). All authors read and approved the final version of the manuscript.

## Data availability

The RNA-seq data have been deposited in NCBI’s Gene Expression Omnibus (Edgar *et al.*, 2002) and are accessible through GEO Series accession number GSE217852 (https://www.ncbi.nlm.nih.gov/geo/query/acc.cgi?acc=GSE217852). The full IDIO+ transcriptome (TSA, transcriptome shotgun assembly) including coding, non-coding and splicing variants, has been deposited at DDBJ/EMBL/GenBank under the accession GKIJ00000000. The version described in this paper is the first version, GKIJ01000000. All other data supporting the findings of this study are available within the paper and within its supplementary data published online.

